# Redox regulation by the CDSP32 thioredoxin of ATP-synthase activity and enzymatic antioxidant network in *Solanum tuberosum*

**DOI:** 10.1101/2024.02.29.582824

**Authors:** Pascal Rey, Patricia Henri, Jean Alric, Laurence Blanchard, Stefania Viola

**Affiliations:** Aix Marseille Univ, CEA, CNRS, BIAM, Photosynthesis & Environment (P&E) Team, Saint Paul-Lez-Durance, F-13115, France; Aix Marseille Univ, CEA, CNRS, BIAM, Molecular and Environmental Microbiology (MEM) Team, Saint Paul-Lez-Durance, F-13115, France

## Abstract

Plant thioredoxins (TRXs) form a complex family involved in numerous metabolic and signalling pathways, such as the regulation of photosynthetic metabolism in relation with light conditions. The atypical CDSP32, chloroplastic drought-induced stress protein of 32 kDa, TRX includes two TRX-fold domains, one of which has an atypical redox-active HCGPC motif, and has been initially reported to participate in responses to oxidative stress as an electron donor to peroxiredoxins and methionine sulfoxide reductases. Here, we further characterized potato lines modified for *CDSP32* expression to clarify the physiological roles of the TRX. Upon high salt treatments, modified lines displayed changes in the abundance and redox status of CDSP32 antioxidant partners, and exhibited sensitivity to NaHCO_3_, but not to NaCl. In non-stressed plants overexpressing CDSP32, a lower abundance of photosystem II PsbO and D1 subunits and ATP-synthase γ subunit was noticed. The CDSP32 co-suppressed line showed altered chlorophyll *a* fluorescence induction and modified regulation of the plastidial ATP-synthase activity during dark/light and light/dark transitions, revealing the involvement of CDSP32 in the control of the photosynthetic machinery. In agreement with the previously reported interaction *in planta* between CDSP32 and the ATP-synthase γ subunit, our data show that CDSP32 participates in the regulation of the transthylakoid membrane potential. Consistently, modeling of protein complex 3-D structure indicates that the CDSP32 TRX constitutes a suitable partner of ATP-synthase γ subunit. We discuss the roles of CDSP32 in chloroplast redox homeostasis through the regulation of both photosynthetic activity and enzymatic antioxidant network.

## Introduction

Redox post-translational modifications (PTM) in proteins lead to switches modifying enzymatic activity, protein conformation or subcellular distribution. In the presence of reactive oxygen species (ROS), cysteine (Cys) can be oxidized to various forms such as sulfenic acid or disulfide bond, while methionine (Met) can be oxidized to Met sulfoxide (Davies, 2005; Delaunay et al. 2002; Tada et al. 2008). Of note, most redox PTMs in Cys and Met are reversible thanks to the action of thiol reductases (TRs), in which thiol deprotonation of a catalytic cysteine generates a nucleophilic reactive thiolate form. By modulating Cys redox status in interacting partners, TRs play key roles in cell redox homeostasis maintenance and ROS-related signalling transduction pathways (Lu and Holmgren, 2014). Plant TRs form complex multigene families including thioredoxins (TRXs) (Meyer et al, 2012; Geigenberger et al. 2017) and glutaredoxins (GRXs), which are closely related to TRXs and use glutathione (GSH) as an electron donor (Rouhier et al. 2008). TRXs are ubiquitous small disulfide reductases carrying a two-Cys active site, generally WCGPC. Cytosolic TRXs get reducing power from NADPH-TRX reductases and are involved in various processes such as mobilization of seed reserves upon germination or biotic stress responses (Sweat and Wolpert, 2007; Hägglund et al. 2016). Plastidial TRX isoforms are mainly supplied with electrons from the PSI/ferredoxin/ferredoxin-TRX-reductase (FTR) pathway and play essential roles, in concert with 2-Cys peroxiredoxins (PRXs), in the regulation of photosynthetic metabolism notably upon dark/light and light/dark transitions (Schürmann and Buchanan, 2008; Courteille et al, 2013; Naranjo et al. 2016; Yoshida et al. 2018; Lampl et al. 2022). Thiol peroxidases such as PRXs also fulfil signalling roles via the control of peroxide concentration or direct thiol oxidation in protein partners (Rhee and Woo, 2011; Cerveau et al. 2016; Liebthal et al. 2018). Other TRX targets like methionine sulfoxide reductases (MSRs) maintain protein redox status thanks to redox-active cysteines (Tarrago et al. 2009; Laugier et al. 2010). By modifying the Cys redox status in a large set of partners (Montrichard et al. 2009), plant TRXs participate in multiple metabolic, developmental and stress-related processes (Gelhaye et al. 2005; Vieira Dos Santos and Rey, 2006; Geigenberger et al. 2017; Montillet et al. 2021).

In addition to canonical TRXs, such as cytosolic h, mitochondrial o and plastidial f, m, x and y isoforms which contain only one TRX-fold domain and a typical active site WCGPC, plants harbour TRXs carrying atypical active site motifs or including other domains (Meyer et al. 2012). Among them, the CDSP32 (Chloroplastic Drought-induced Stress Protein of 32 kDa) TRX, initially identified in potato plants subjected to water deficit (Rey et al. 1998), is induced by various environmental constraints (Pruvot et al. 1996; Rey et al. 1998; Broin et al. 2000). The protein is composed of two TRX-fold domains with the N-terminal one having no redox-active motif, but instead a SXXS motif, and the C-terminal one displaying an atypical redox-active HCGPC motif (Rey et al. 1998). The involvement of CDSP32 in responses to oxidative stress was established based on the phenotype of transgenic lines (Broin et al. 2002; 2003) and on its capacity to reduce various PRXs and MSRs (Rey et al. 2005; Vieira Dos Santos et al. 2007; Tarrago et al. 2010). Accordingly, ectopic expression of mulberry and cotton *CDSP32* genes in Arabidopsis was found to trigger recovery following a drought period (Sun et al. 2020) and to confer tolerance to osmotic or oxidative stresses (Elasad et al. 2020).

Besides this role in stress response, CDSP32 could participate in photosynthesis regulation. Indeed, in CDSP32-overexpressing tobacco plants, Zhang et al. (2020) noticed higher chlorophyll (Chl) content together with a reduced Chl a/b ratio, and better maintenance of photosynthetic activity upon cadmium treatment, compared to wild-type (WT). They hypothesized that the TRX regulates photosynthetic cyclic electron flow and energy dissipation allowing photoprotection in stress conditions (Zhang et al. 2021). Most importantly, by performing redox proteomics during a dark/light transition in tobacco, Zimmer et al. (2021) revealed that the redox status of both TRX f and CDSP32 is dependent on the rate of linear electron flow, and proposed that the two TRXs act in concert in the regulation of Calvin-Benson cycle activity.

Here, we further characterized potato lines modified for *CDSP32* expression to better delineate the physiological roles of the TRX. Upon high salt treatments, modified lines displayed substantial changes in the abundance and redox status of CDSP32 partners, and exhibited sensitivity to NaHCO_3_, but not to NaCl. In control conditions, changes in the abundance of some photosystem II (PSII) subunits and in chlorophyll content were noticed in lines overexpressing CDSP32, whereas the line lacking the TRX displayed modified regulation of the chloroplast ATP-synthase activity during dark/light and light/dark transitions, in agreement with the identification of the ATP-synthase γ subunit as an *in planta* CDSP32 partner (Rey et al. 2005). Based on these results, we discuss the roles of CDSP32 in the regulation of chloroplast redox homeostasis through its participation in both control of photosynthetic activity and antioxidant network.

## Materials and methods

### Sequence analysis and 3D structure modelling

Sequences of CDSP32 TRXs and related proteins and of ATP-synthase γ subunit among plant, algae and bacteria phyla were found by searching databases and performing BLAST analyses at NCBI: https://blast.ncbi.nlm.nih.gov/Blast.cgi. Prediction of transit peptide was performed at DTU (Technical University of Denmark): https://services.healthtech.dtu.dk/services/TargetP-2.0/. Multiple sequence alignments were carried out using Clustal Omega at EMBL-EBI: https://www.ebi.ac.uk/Tools/msa/clustalo/. Analysis of conserved protein domain family was performed at NCBI: https://www.ncbi.nlm.nih.gov/Structure/cdd/wrpsb.cgi (Wang et al. 2023). The 3D structures of *S. tuberosum* CDSP32 and TRX f were predicted using AlphaFold2 via the software ColabFold (Jumper et al. 2021; Mirdita et al. 2022). The 3D structures of the *S. tuberosum* ATP synthase γ subunit and the complexes between the γ subunit and CDSP32 and TRX f were predicted using a local AlphaFold2 installation (version 2.3.1) (Jumper et al. 2021). All parameters were kept to default values. The quality of each model, presented in supplementary figures, was evaluated using the confidence measures pLDDT (predicted local distance difference test) that indicates the local accuracy, and PAE (predicted aligned error) that assesses the packing between domains or protein chains. The 3D structure of *Spinacia oleracea* ATP-synthase and of its γ subunit was obtained from the protein data bank (https://www.rcsb.org, PDB 6FKF). Three-dimensional structure or model images (rank_1 model for each AlphaFold2 predicted 3D structure) were generated using PyMOL (PyMOL Molecular Graphics System, Version 2.0 Schrödinger, LLC). The 3D modelled structures of proteins or complexes do not include transit peptides, therefore the sequence numbering of all proteins starts after the adressing peptides.

### Plant material and growth conditions

Potato (*Solanum tuberosum* L. cv Désirée) lines were propagated *in vitro* on Murashige and Skoog medium, and transferred for growth *in vivo* in a phytotron under LED light (VEGELED Horticulture Floodlights Apollo LL300, Colasse SA, Belgium) (PPFD of 185 µmol.m^-2^.s^-1^, 12 h day/12 h night, 23/19°C) in the “Phytotec” platform (CEA, DRF, BIAM). In addition to WT, three transgenic lines co-suppressed for the expression of the potato *CDSP32* gene (termed D4), overexpressing it (D10) or overexpressing in the WT background a mutated form of the TRX, where the catalytic Cys is replaced by a Ser (DM19), were used (Broin et al. 2002; Rey et al. 2005). Following transfer on soil, control plants were watered with water the first three days, and then with nutritive solution (Coic and Lesaint, 1971). For salt treatments, the nutritive solution was supplemented with of 0.125 M NaCl or 0.1 M NaHCO_3_. The treatments were applied for 21 days. At this stage, plant height was measured and aerial parts weighed, and samples for chlorophyll determination and protein analyses collected.

### Protein extraction and content determination

Pieces from the terminal leaflet of young expanded leaves were blended in liquid N_2_, and the powder was resuspended in 50 mM Tris-HCl, pH 8.0, 1 mM phenylmethylsulfonyl fluoride, and 50 mM β-mercaptoethanol. Following centrifugation (14,000 g, 4°C, 20 min), soluble proteins were recovered by precipitation of the supernatant using two acetone volumes and stored at −20°C. The pellet, containing thylakoids, was resuspended in 50 mM Tris-HCl pH 8.0, 1% SDS, agitated for 2 h at 4°C, then centrifuged (14,000 *g*, 4°C, 20 min). Membrane proteins were precipitated at −20°C by the addition of four volumes of acetone to the supernatant. Protein concentration was quantified using the “Protein Quantification BCA Assay” kit (Pierce BCA Protein Assay Kit, Thermo Fisher Scientific).

### Western blot analysis

Proteins were separated by SDS-PAGE generally in 13%, or when needed in 8, 10, 11 or 15%, acrylamide gels (Laemmli, 1970) in reducing conditions (0.1 M DTT in solubilization buffer) and electroblotted onto 0.45 µM nitrocellulose (Pall Gelman Sciences). Membranes were stained with Ponceau red to ensure homogenous and proper transfer. The list of primary antibodies raised in rabbit used in this work is presented in Table S1. Membranes could be subsequently subjected to two distinct revelation procedures. In the first procedure, bound antibodies were detected using a goat anti-rabbit secondary antibody coupled to a fluorescent molecule (Alexa Fluor 680, Invitrogen) diluted 1:10,000 using the “Odyssey Infrared Imager” at 680 nm (Licor, Lincoln, NE, USA). Quantification of band intensity was performed using the software associated with the imager. Then the membrane was incubated with another primary serum and revealed using an anti-rabbit immunoglobulin G coupled to alkaline phosphatase (Sigma). Data were acquired from three or four independent experiments and representative immunoblots are shown. Quantification of band intensities were obtained from biological replicates yielding consistent results.

### Chlorophyll content determination

Leaf disks of 0.6 cm diameter were collected on terminal leaflets of young expanded leaves, stored at – 80°C before blending in 80% acetone. Following shaking, overnight storage in the dark at 4°C and centrifugation (14,000 g, 20 min), the content in chlorophylls *a* and *b* was measured by spectrophotometry at 647 and 663 nm, and calculated according to Lichtenthaler (1987).

### Fluorescence measurements

Chlorophyll fluorescence was measured on leaf fragments using a laboratory-built fluorescence camera. Leaf fragments were harvested from plants incubated in darkness overnight (>12h), and kept on wet Whatman paper in darkness until measurements were performed. Each biological replicate consisted in one leaf fragment harvested from a different plant. Maximal PSII quantum yield [Fv/Fm = (Fm–Fo)/Fm] was measured applying a multi-turnover saturating light pulse (200 ms, 3,000 µmol photons m^-2^ s^-1^) to dark-adapted leaves. Fluorescence induction curves were measured during 4 minutes of actinic light at the indicated intensities, and Fm’ levels after multi-turnover saturating pulses applied during the actinic illumination. PSII quantum yield [Φ_PSII_ = (Fm’-F)/Fm’] was calculated with F and Fm’ values measured, respectively, right before and after the first and last saturating pulses of the sequence. Non-Photochemical Quenching (NPQ) was calculated as (Fm-Fm’)/Fm’. Actinic light and saturating pulses were provided by orange-red LEDs, measuring pulses (740 µs) by blue LEDs.

### ElectroChromic Shift (ECS) measurements

ECS measurements were performed with a laboratory-built Joliot-Type Spectrophotometer (JTS). Short (15 µs) measuring pulses were provided by a sun-like LED (COB V6HD THRIVE WHT SQ 6500K, Bridgelux) filtered via a 520 nm interference filter (Edmund Optics, 10 nm FWHM). Actinic light was provided by orange-red LEDs. Single-turnover saturating flashes (700 nm, duration 6 ns) were provided by a pulsed Nd:YAG laser pumping an optical parametric oscillator (Surelite II, Continuum). For sub-saturating flashes, the laser light was attenuated using a metal grid. Measurements were performed on intact leaves detached right before measuring from plants incubated in darkness overnight (>12h). Each biological replicate consisted in one leaf detached from a different plant.

### Electron Transport Rate (ETR) measurements

Dark-adapted leaves were illuminated with the indicated intensities of actinic light for 4 minutes. The PSII+PSI antenna size and total steady-state ETR were calculated from the slopes of the absorption changes at the onset and offset of the actinic illumination, respectively (for details see Mathiot and Alric, 2021). The absorption changes induced by a single-turnover saturating flash, inducing one charge separation per photosystem (PSII+PSI) were used for normalisation.

### Redox-dependent regulation of the ATP-synthase activity

Activation: absorption changes induced by one sub-saturating flash were recorded in dark-adapted leaves and 2 minutes after a train of n sub-saturating pre-flashes fired at 20 Hz, with n = 20, 50, 100, 200 and 300. For each leaf, the loss in amplitude of the slow ECS decay phase induced by n pre-flashes (-A_n_) was calculated with respect to the amplitude of the slow ECS decay phase in the dark-adapted leaf (A_Dark_), setting the amplitude of the slow phase after 300 pre-flashes as the zero value. The number of charge separations per PSII+PSI induced by n pre-flashes was calculated by multiplying n for the ratio between the absorption changes induced by a sub-saturating and a saturating flash.

De-activation in darkness: absorption changes induced by one sub-saturating single-turnover flash were recorded in dark-adapted leaves and at different times t after a train of 300 sub-saturating pre-flashes fired at 20 Hz, with t = 2, 4, 6, 8, 10, 15 and 20 minutes. For each leaf, the recovery in amplitude of the slow ECS decay phase at a time t after the 300 pre-flashes (A_t_) was calculated with respect to the amplitude of the slow ECS decay phase in the dark-adapted leaf (A_Dark_), setting the amplitude of the slow phase at t = 2 min as the zero value.

For each curve, the –A_n_/A_Dark_ (activation measurements) and A_t_/A_Dark_ (de-activation measurements) values calculated for the absorption changes measured at 395, 595, 789 and 985 ms after the flash were averaged.

## Results

### CDSP32 presence in the green lineage

We first performed an extensive search of databases to investigate the presence of CDSP32-related proteins in living organisms. We retained proteins displaying only the two very specific features of the TRX, *i.e.* two TRX-fold domains and an atypical active site motif, HCGPC. Indeed, based on the classification of conserved protein domains (Wang et al. 2023), the two N-ter and C-ter domains, which contain SXXS and CXXC motifs, respectively, belong to the cl00388 thioredoxin-like superfamily, which is subdivided in many classes corresponding to various types of thiol reductases. CDSP32-related proteins form a unique subfamily, cd02985, termed TRX_CDSP32. All Dicotyledon and Monocotyledon species analysed harbour *CDSP32* genes encoding very well conserved proteins, except the transit peptide, as shown by multiple sequence alignment (Fig. S1). For the species selected, N- and C-terminal CDSP32 domains exhibit *ca.* 40% and more than 70% identity, respectively. We particularly noticed the presence of a conserved SXXS motif in the N-ter domain at the active site position in the C-ter domain (Rey et al. 1998), and very high identity in the sequence surrounding the active site. This highly conserved sequence constitutes a typical CDSP32 19-residue motif, VLDVGLK<u>HCGPC</u>VKVYPTV, not found in plant canonical TRXs. When extending the homology search to other plant groups (Fig. 1), we observed that the protein is well-conserved in primitive Angiosperms such as *Amborella trichopoda*, found only in one class of Gymnosperms, Pinales, and present in several classes of Pteridophytes. The CDSP32 homologues from this latter group also possess the typical 19-residue motif, which differs by two residues in Lycopodiopsida. In Bryophytes, related proteins are present in Mosses and Liverworts, but display sequence divergence in the active site (NCGPC and SCGPC in *Physcomitrium patens* and *Marchantia polymorpha*, respectively) and in the surrounding sequence. Alignments of sequences from non-Algae photosynthetic organisms revealed more than 40% identity of the C-terminal domain (Fig. S2). In green Algae, the sequence identity was much lower (Fig. S3). The CDSP32 homologous proteins identified in Charophytes, such as *C. braunii*, harbour active sites containing two Cys residues, but no His preceding the catalytic Cys. In Chlorophytes, very poor conservation of the typical CDSP32 sequence motif was observed, since the potential active site carries only one Cys, for example SAGPC in *Chamydomonas reinhardtii* (Fig. 1).

**Figure 1.**
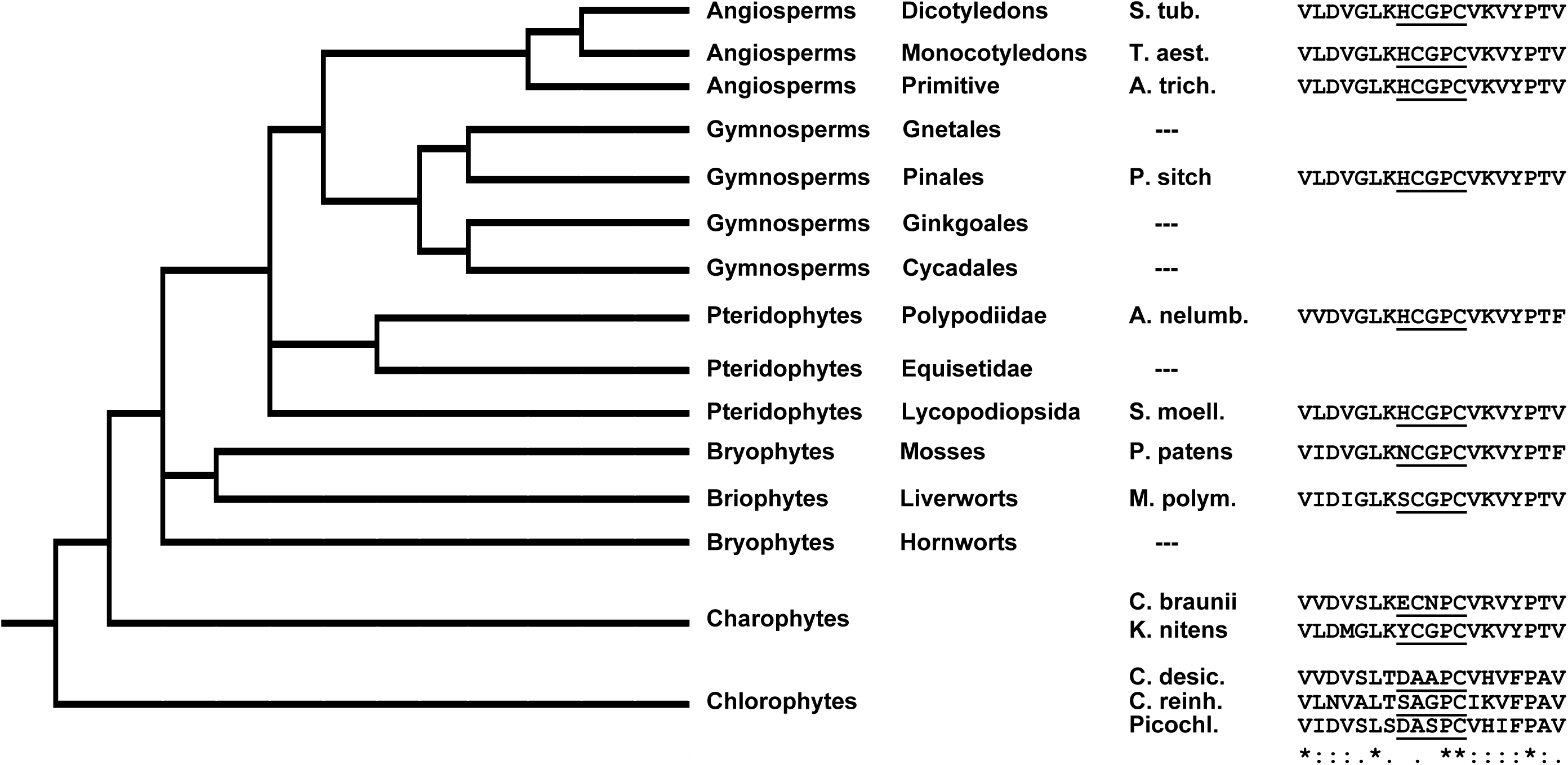
Presence of the CDSP32 thioredoxin and related proteins among the green lineage. CDSP32 sequences were analyzed from several species of each main representative group of the green lineage. Typical 19-residue motifs surrounding the active site, which is underlined, are shown. Species and accession numbers : S. tub., *Solanum tuberosum* (NP_001305492.1); T. aest.; *Triticum aestivum* (XP_044421501.1); A. trich., *Amborella trichopoda* (XP_006828837.1); P. sitch., *Picea sitchensis*; (ABK25354.1); A. nelumb., *Adiantum nelumboides* (L7F22_035310); S. moell., *Selaginella moellendorffii* (XP_002989497.2); P. patens, *Physcomitrium patens* (XP_024392553.1); M. polym., *Marchantia polymorpha* (PTQ43510.1); C. braunii, *Chara braunii* (CBR_g26413); K. nitens, *Klebsormidium nitens* (KFL_004880100); C. desic., *Chlorella desiccata* (KSW81_007621); C. reinh., *Chlamydomonas reinhardtii* (CHLRE_14g624201v5); Picochl*., Picochlorum* sp. BPE23 (M9435_003533).

In conclusion, the CDSP32 TRX, although possibly absent from some classes of Gymnosperms and Pteridophytes, is particularly well conserved in vascular plants. The presence of related proteins in Bryophytes and green Algae indicates that this TRX type is represented in most photosynthetic eukaryotes of the green lineage, although the typical TRX activity might not yet been acquired in Chlorophytes. No homologous proteins were identified in Cyanobacteria, but very surprisingly, some bacterial sequences encoding hypothetical proteins with two TRX-fold domains share high identity with Angiosperm CDSP32 (Fig. S4). For example, hypothetical proteins from *Nitriliruptoraceae bacterium* and *Enterobacter hormaechei* possess more than 70% identity with mature *S. tuberosum* CDSP32 and contain the 19-residue motif, except an active site RCGPC in *Enterobacter*, raising the question of the origin of the gene in these Prokaryotes.

### CDSP32 structure features

We then investigated whether the CDSP32 sequence features confer specific structural and electrostatic characteristics to the TRX notably compared to canonical plastidial ones such as TRX f. The 3D structure of *S. tuberosum* CDSP32 predicted by AlphaFold2 shows two TRX-fold domains as expected (TRX_D1 and TRX_D2), each with four α helices surrounding four β strands, separated by a 10-residue flexible region (Fig. 2A, Fig. S5). The AlphaFold2 structure of *S. tuberosum* TRX f shows a single TRX-fold domain that perfectly superimposes on the TRX_D2 domain of CDSP32 including the same position of the redox site motif (H/WCGPC) (Fig. 2A, Fig. S6). Although these TRXs show the same folding, they present different electostatic properties (Fig. 2B) that can play a crucial role in distinct substrate specificity via electrostatic complementarity with their partners as shown for *Arabidopsis thaliana* TRXs isoforms (Bodnar et al. 2023). Of note, the two TRX-fold domains of CDSP32 (that share 26.8 % identity) also perfectly superimpose but show different electrostatic properties (Fig. 2B, Fig. S5).

**Figure 2.**
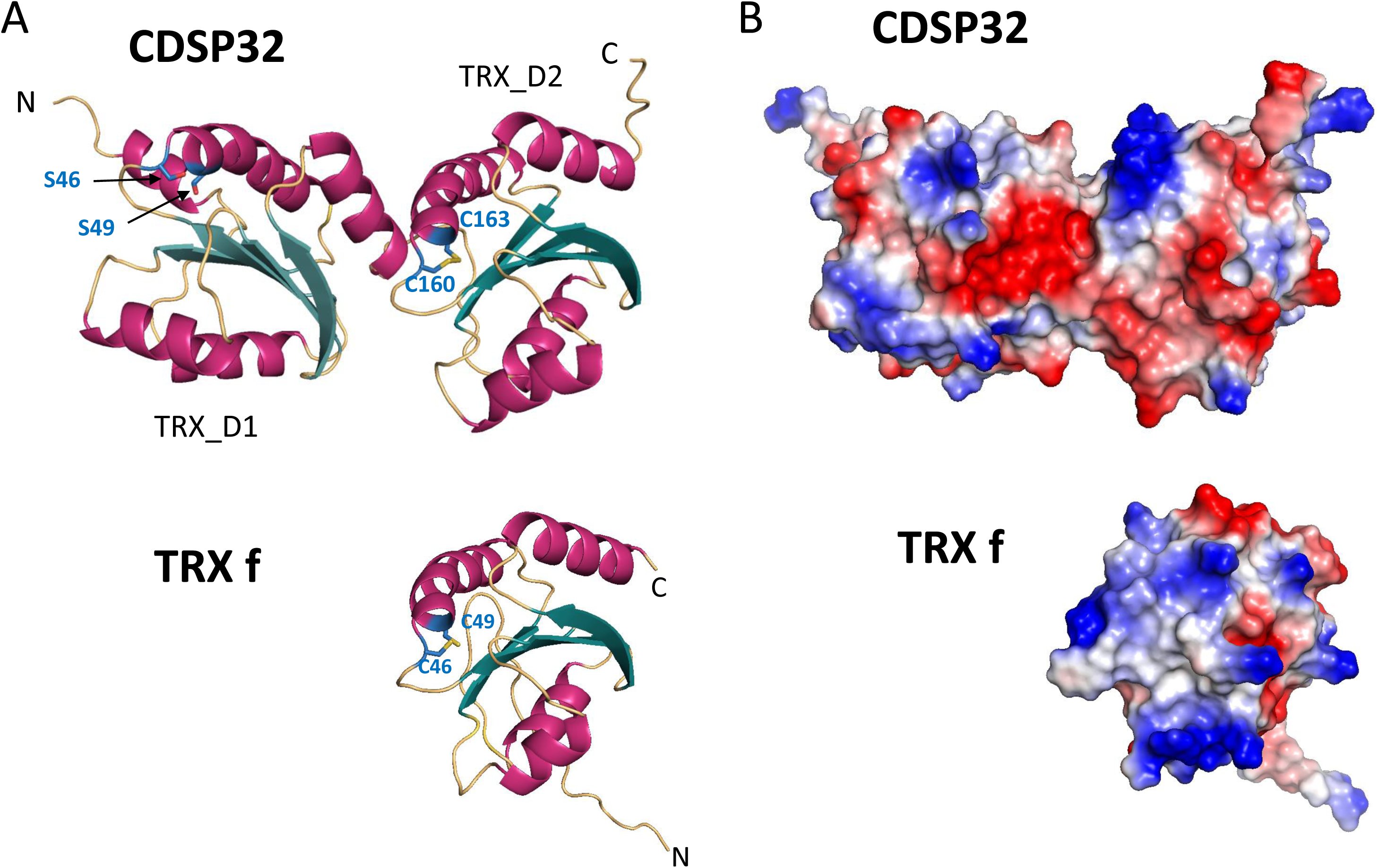
AlphaFold2 structure of *S. tuberosum* CDSP32 (top) and TRX f (bottom). (A) The 3D structures are colored according to secondary structure elements (α-helices in pink and β-strands in teal blue). CDSP32 presents two TRX-fold domains (D1 and D2) in tandem, and TRX f a single TRX-fold domain. The Cys residues of the active motif and the Ser residues of the corresponding motif in CDSP32 TRX_D1 are shown in blue, (B) Surface representation with the same orientation as in (A) displaying atoms colored-coded according to the surface electrostatic potential from red (negative) to blue (positive). Different electrostatic properties are observed between CDSP32 and TRX f.

### Growth characteristics and chlorophyll content in control conditions

We previously reported the generation of potato lines either lacking the CDSP32 TRX, or overexpressing the WT protein or the Cys-216-Ser active site mutated form (Broin et al. 2002; Rey et al. 2005). In these first assays, we did not notice any obvious growth or development phenotype for these lines in the absence of environmental constraints. As these assays were performed on a limited number of plants and under a different light source (HQI-sodium lamps) than the one used in this work (LED lamps), we first carried out a deeper phenotype characterization in control conditions. WT and transgenic lines displayed similar growth and development features after 3.5 weeks following the transfer from *in vitro* tubes (Fig. 3A). All lines exhibited stem heights in the range of 21 cm and weights of aerial parts close to 30 g (Fig. 3B, C). Based on the somewhat paler leaf colour in plants expressing mutated CDSP32, we determined the chlorophyll content of young expanded leaves in the upper crown (Fig. 3D). While WT, co-suppressed (D4) and CDSP32-overexpressing (D10) plants displayed contents between 36 and 38 µg. cm^-2^, a significantly lower content (less than 33 µg. cm^-2^) was measured in plants overexpressing the mutated TRX (DM19). No difference in the chl *a*/*b* ratio was observed, indicating that both chlorophyll types were similarly affected in this line (data not shown).

**Figure 3.**
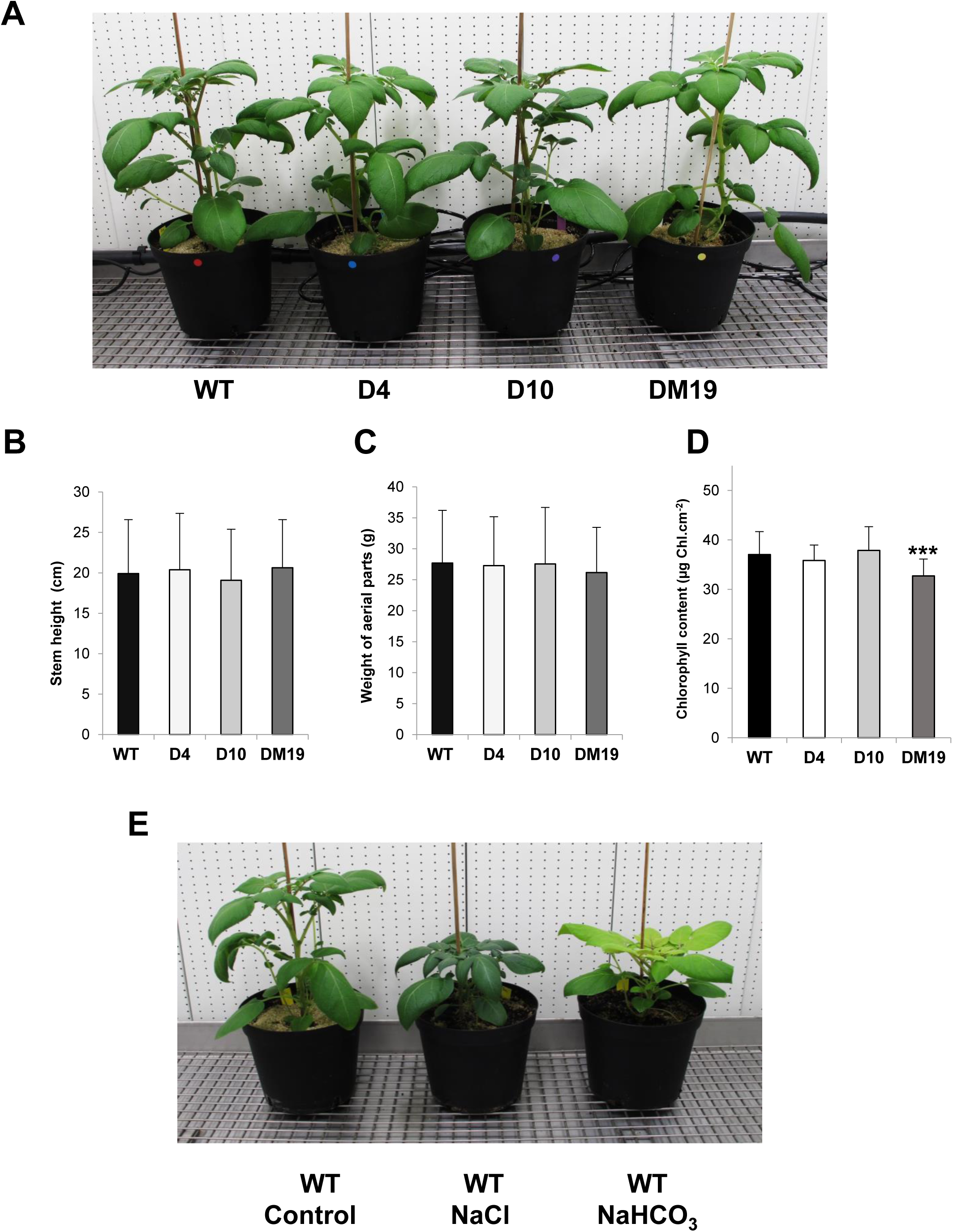
Growth of potato plants modified in *CDSP32* expression upon standard or high salinity conditions. **A.** Three-week old plants grown on soil in a phytotron under standard watering conditions. **B.** Histogram representation of the stem height of 24-day old plants grown in standard conditions. **C.** Histogram representation of the weight of aerial parts of 24-day old plants grown in standard conditions. **D.** Histogram representation of the chlorophyll content of young expanded leaves from the upper crown of 24-day old plants grown in standard conditions. **E.** Three-week old WT plants watered with standard nutritive solution or solutions containing 0.125 M NaCl or 0.1 M NaHCO_3_. WT, wild type; D4, line co-suppressed for CDSP32; D10, line overexpressing CDSP32; DM19, line overexpressing CDSP32 active site mutant. Growth data are means ± SD of six average values originating from independent experiments (four plants for each genotype). Chlorophyll data are means ± SD from 35 independent measurements. ***, significantly different from the WT value with P < 0.001 (t-test).

### Growth characteristics upon high salt treatments

*CDSP32* gene expression is upregulated in WT potato plants upon severe osmotic stress conditions such as drought or exposure to high NaCl concentration (0.3 M) (Pruvot et al. 1996). Here, we subjected transgenic potato lines to a lower NaCl concentration (0.125 M) in watering solution for three weeks. Upon NaCl exposure, the stem height was strongly reduced in all lines (12 cm *vs* 21 cm in control conditions (Figs. 3E, S7A, B). The treatment also lead to a dramatic decrease in the weight of aerial parts to *ca.* 17 g (against 30 g in control conditions) in the four potato lines, plants overexpressing WT CDSP32 having a slightly higher weight (Fig. S7C). We noticed that NaCl-treated plants were greener than control ones (Fig. 3E). Consistently, we measured a chlorophyll content higher than 45 µg.cm^-2^ in the same range in all lines (Fig. S7D). This increased level is likely due to the leaf size reduction and thickness increase induced by the NaCl treatment, resulting in higher chloroplast density per leaf surface area (Munns and Tester, 2008). Altogether, these data show that the phenotype of potato plants treated with 0.125 M NaCl is unaffected by modifications of *CDSP32* expression.

Since some CDSP32 targets like MSRBs are involved in responses to saline-alkaline conditions (Sun et al. 2016) and *CDSP32* expression is impaired in such stress conditions (Huihui et al. 2020), we exposed plants to 0.1 M NaHCO_3_ for three weeks. Similarly to NaCl treatment, watering with NaHCO_3_ resulted in a strong decrease in plant size for all lines (Fig. 3E and 4A) with a stem height in the range of 12 cm (Fig. 4B). The weight of aerial parts was also strongly reduced upon salt treatment. Interestingly, WT and CDSP32-overexpressing plants exhibited weights of around 15 and 18 g, respectively, while a substantially lower value, 11 g, was recorded for plants overexpressing the mutated form (Fig. 4C). Exposure to NaHCO_3_, in contrast to NaCl, did not affect leaf expansion (Fig 3E). The four potato lines watered with NaHCO_3_ displayed pale green to yellow leaves, particularly the youngest ones (Fig. 4A). Chlorophyll content measurements revealed that WT plants were less affected (15.5 µg.cm^-2^) compared to CDSP32-modified plants, particularly those overexpressing mutated CDSP32 (*ca.* 9 µg.cm^-2^) (Fig. 4D). Of note, another line overexpressing mutated CDSP32 termed DM15 (Rey et al. 2005) also displayed reduced growth and paler leaves compared to WT (Fig. S8A, B). In other respects, we noticed that plants treated with NaHCO_3_ displayed severe necrotic lesions in older leaves and that WT plants exhibited less damaged leaves compared to CDSP32-modified lines (Fig. S8C, D). Taken together, these data reveal that plants modified for CDSP32 expression, especially those overexpressing the mutated form, are more sensitive than WT to NaHCO_3_ exposure.

**Figure 4.**
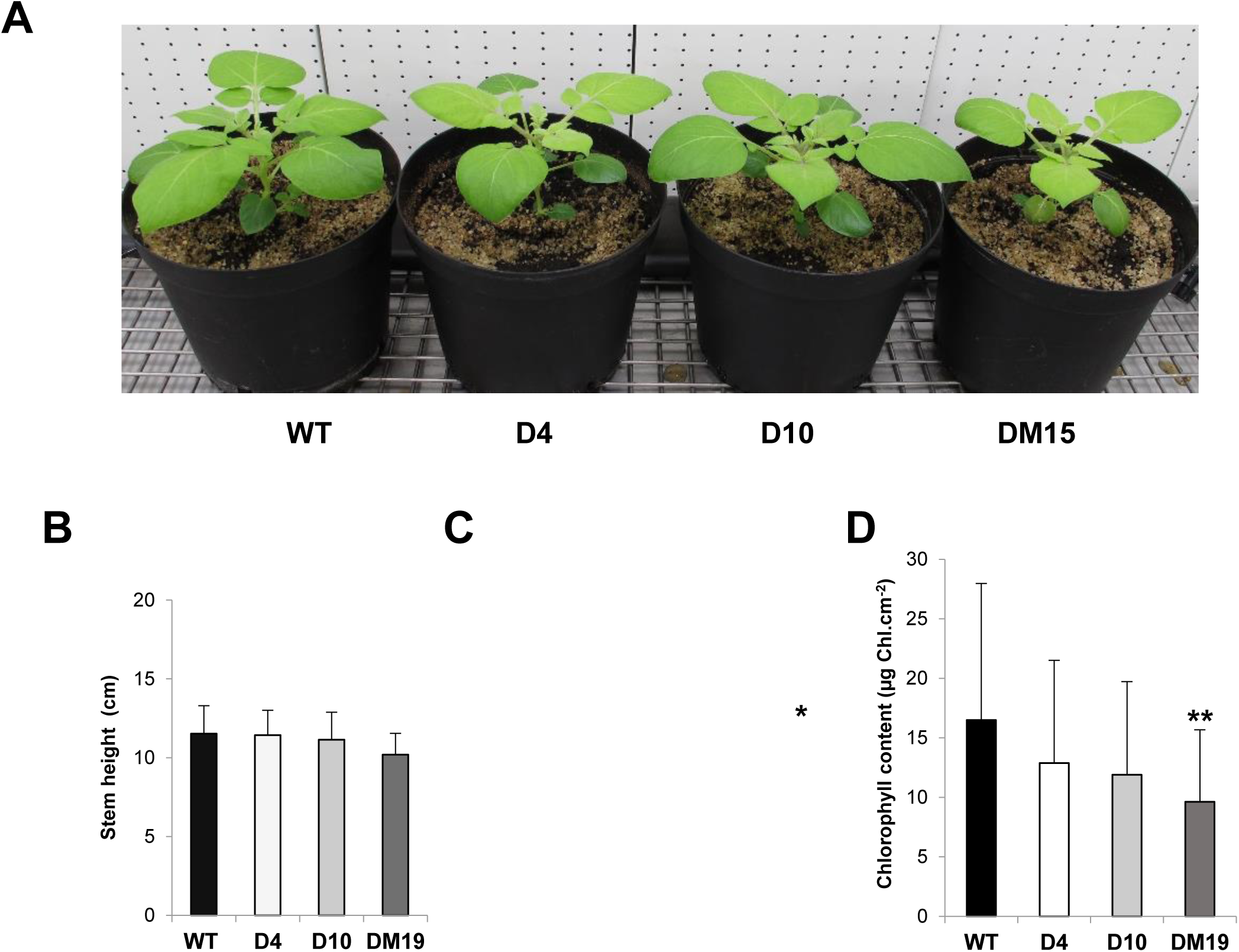
Growth of potato plants modified in *CDSP32* expression upon NaHCO_3_ exposure. **A.** Three-week old plants grown on soil in a phytotron were watered with a nutritive solution containing 0.1 M NaHCO_3_. **B.** Histogram representation of the stem height of 24-day old plants exposed to NaHCO_3_. **C.** Histogram representation of the weight of aerial parts of 24-day old plants exposed to NaHCO_3_. **D.** Histogram representation of the chlorophyll content of young expanded leaves from the upper crown of 24-day old plants exposed to NaHCO_3_. WT, wild type; D4, line co-suppressed for CDSP32; D10, line overexpressing CDSP32; DM19, line overexpressing CDSP32 active site mutant. Growth data are means ± SD of six average values originating from independent experiments (four plants for each genotype). Chlorophyll data are means ± SD from 35 independent measurements. * and **, significantly different from the WT value with P < 0.05 and P < 0.01, respectively (t-test).

### Abundance of CDSP32 partners

We performed Western blot analyses to investigate the abundance of several thiol reductases, most being known to interact with CDSP32. First, we reassessed the abundance of the CDSP32 TRX in the four potato lines grown in control conditions (Fig. 5A). While the protein was not detected in the D4 co-suppressed line, much higher abundances were observed compared to WT in over-expressing lines, with 5- and 8-fold more intense bands in plants expressing WT or mutated forms, D10 and DM19, respectively. In agreement with previous reports (Pruvot et al. 1996), NaCl treatment lead to a noticeable increase in the abundance of the TRX in WT plants. Upon watering with NaHCO_3_ (Fig. 5A), no change in protein abundance was noticed in WT plants, and decreased amounts were observed in overexpressing lines compared to control conditions.

**Figure 5.**
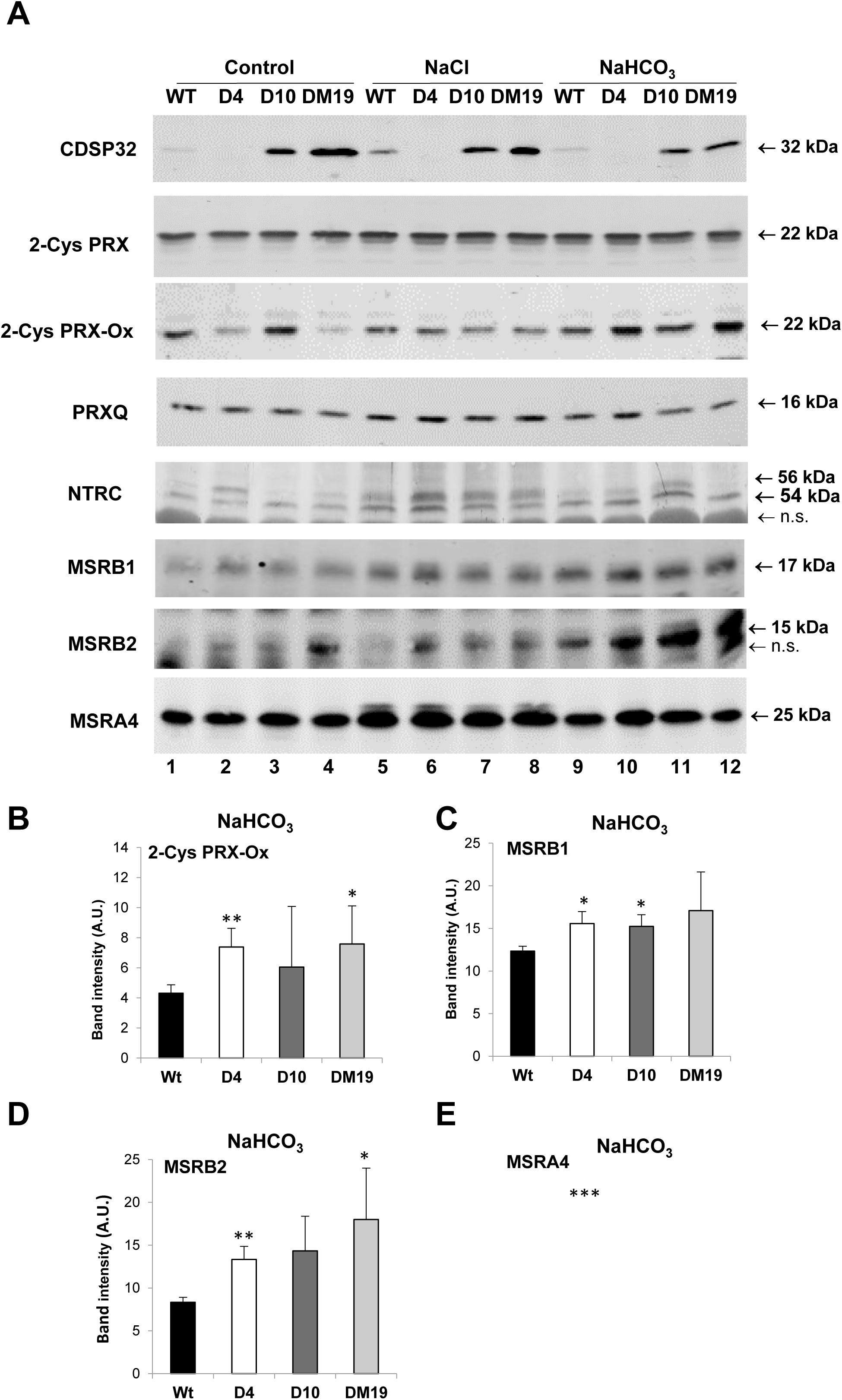
Abundance of CDSP32 and various CDSP32-related plastidial thiol reductases in potato plants modified in *CDSP32* expression grown in standard conditions or exposed to high salinity. **A.** Western blot analysis of the abundance of CDSP32, 2-Cys PRX, overoxidized 2-CysPRX (2-Cys PRX-Ox), PRXQ, NTRC, MSRB1, MSRB2 and MSRA4 in plants grown in standard conditions or exposed to high salinity treatments, 0.125 M NaCl or 0.1 M NaHCO_3_. **B.** Histogram representation showing the abundance of 2-Cys PRX Ox and of MSRs in plants exposed to NaHCO_3_. Data are means ± SD of four values originating from independent experiments. *, * and ***, significantly different from the WT value with P < 0.05, P < 0.01 and P < 0.001, respectively (t-test). WT, wild type; D4, line co-suppressed for CDSP32; D10, line overexpressing CDSP32; DM19, line overexpressing CDSP32 active site mutant.

We then analysed the abundance of known partners of CDSP32, namely PRXs and MSRs (Fig. 5A). No change in 2-Cys PRX abundance was noticed in control conditions and upon salt treatments depending on *CDSP32* expression level. In control conditions, as previously reported (Cerveau et al. 2016), the 2-Cys PRX over-oxidized form was less and more abundant in co-suppressed and WT-overexpressing lines, respectively. Interestingly, this form involved in redox signalling (Rey et al. 2007) was much less abundant in plants over-expressing inactive TRX than in those expressing the WT form (Fig. 5A, lanes 3-4). Exposure to salts resulted in substantial changes in the level of over-oxidized PRX. Upon NaCl treatment, almost no difference was noticed in the abundance of this form among the four genotypes. In plants watered with NaHCO_3_, higher abundances of over-oxidized PRX were revealed in co-suppressed plants and those over-expressing the mutated TRX (Fig. 5A, B) in sharp contrast with control conditions. With regard to PRXQ, no noticeable variation was observed in the four lines either in control conditions or upon salt treatments (Fig. 5A). Plastidial NADPH-TRX-reductase C, NTRC, is a main reducer of 2-Cys PRX (Pulido et al. 2010), and involved in response to high salt (Serrato et al. 2004). In control conditions, a band at *ca.* 54 kD was revealed in WT by the serum raised against Arabidopsis NTRC, with another upper band at *ca.* 56 kDa. The latter, more abundant in all lines subjected to NaCl treatment, likely corresponds to another redox form, which is clearly detected in non-reducing conditions (Chae et al. 2013). CDSP32 efficiently reduces MSRB1 (Tarrago et al. 2010) and exhibits a weak reducing capacity towards MSRB2 (Vieira Dos Santos et al. 2007). No substantial and reliable change in MSRB1 abundance was observed depending on the potato genotype in control or NaCl conditions (Fig. 5A). In plants watered with NaHCO_3_, a higher MSRB1 abundance was observed in the three transgenic lines compared to WT (Fig. 5A, C). Plants overexpressing mutated CDSP32 displayed a higher MSRB2 abundance in control conditions, and upon salt treatments the three CDSP32-modified lines were also characterized by more elevated MSRB2 protein levels, particularly when watered with NaHCO_3_ (Fig. 5A, D). Finally, we did not observe any substantial variation in MSRA4 amount, except in co-suppressed plants treated with NaHCO_3_, which showed a higher level compared to other lines (Fig. 5A, E). All these data indicate that modifying *CDSP32* expression level substantially affects the redox status or abundance of plastidial PRXs and MSRs notably in NaHCO_3_ salt-stress conditions.

### Abundance of proteins involved in maintenance of redox homeostasis

We investigated the abundance of several ROS-scavenging enzymes, since CDSP32 interacts with 2-Cys PRX, a main player in the maintenance of plastidial ROS homeostasis. Several isoforms of ascorbate peroxidase (APX), which catalyse the H_2_O_2_-dependent oxidation of ascorbate, are present in plant cells, and the serum revealed stromal, peroxisomal and cytosolic APXs in potato extracts (Fig. 6A). In control condition, no change was observed depending on genotype, except somewhat higher levels of peroxisomal and stromal isoforms in D10 plants. Upon NaCl treatment, no substantial change was observed compared to control conditions. In contrast, watering with NaHCO_3_ resulted in a strong decrease in the level of cytosolic APX in all lines (Fig. 6A). Catalase (CAT) is a major peroxisomal enzyme catalysing the decomposition of H_2_O_2_ to water and oxygen. No significant variation in CAT abundance was observed depending on genotype in control and NaCl conditions. In plants watered with NaHCO_3_, a lower protein level was noticed in the three transgenic lines compared to WT (Fig. 6A, B). Superoxide dismutases (SOD) are metalloproteins catalysing dismutation of superoxide anion to oxygen and H_2_O_2_. In control conditions and upon NaCl treatment, the abundance of the plastidial Cu-Zn isoform was similar in all lines. In plants watered with NaHCO_3_, we noticed a significantly higher Cu-Zn SOD amount (by more than 40%) in plants over-expressing WT CDSP32 compared to WT (Fig. 6A, C).

**Figure 6.**
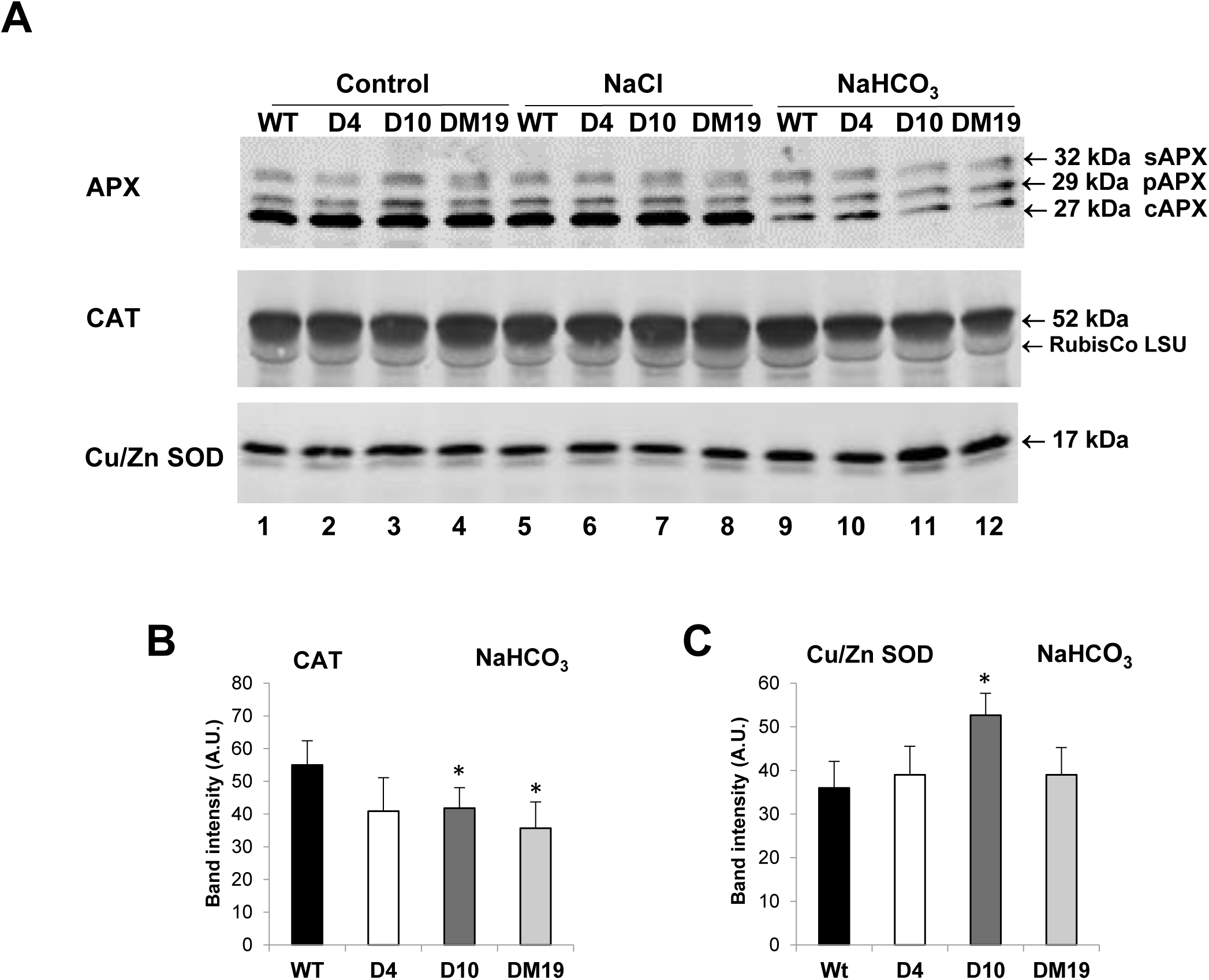
Abundance of ROS-scavenging enzymes in potato plants modified in *CDSP32* expression grown in standard conditions or exposed to high salinity. **A.** Western blot analysis of the abundance of APX, CAT and Cu/Zn SOD in plants grown in standard conditions or exposed to high salinity treatments, 0.125 M NaCl or 0.1 M NaHCO_3_. **B.** Histogram representation showing the abundance of CAT and Cu/ZN SOD in plants exposed to NaHCO_3_. Data are means ± SD of four values originating from independent experiments. *, significantly different from the WT value with P < 0.05 (t-test). WT, wild type; D4, line co-suppressed for CDSP32; D10, line overexpressing CDSP32; DM19, line overexpressing CDSP32 active site mutant.

### Abundance of photosynthetic components and photoprotective proteins

We analysed the abundance of various photosynthetic components, starting with enzymes participating in the Calvin-Benson cycle (CBC), whose activity is redox regulated by TRXs. No genotype-dependent variation in large RuBisCO subunit abundance was observed in plants grown in control conditions or treated with NaHCO_3_, while somewhat higher levels were observed upon NaCl treatment (Fig. 7A). Phosphoribulokinase (PRK) allows the regeneration of ribulose 1,5-bisphosphate, the CO_2_-acceptor substrate in CBC. No substantial variation was noticed in PRK abundance as a function of growth condition or of genotype (Fig. 7A).

**Figure 7.**
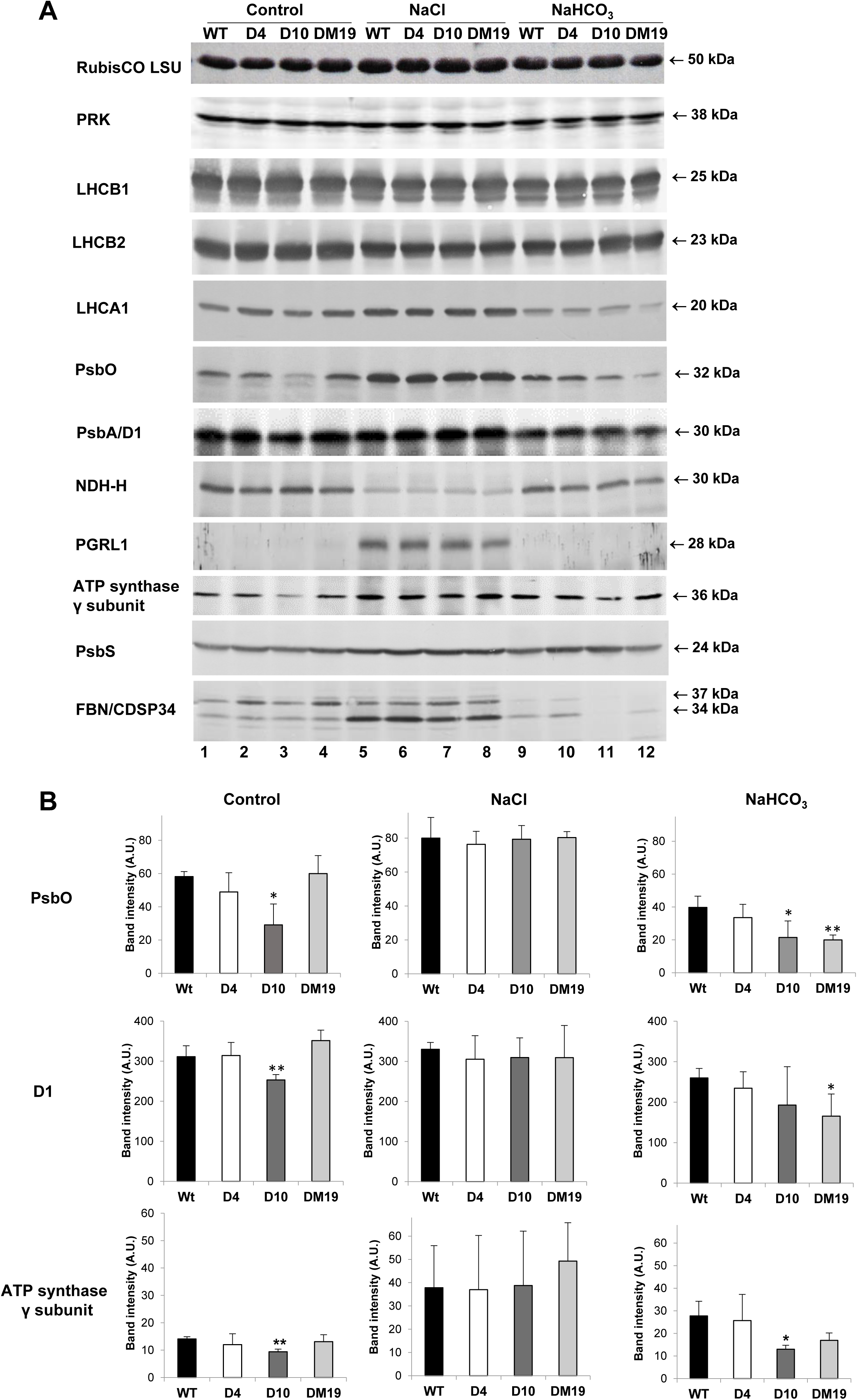
Abundance of photosynthetic and photoprotective components in potato plants modified in *CDSP32* expression grown in standard conditions or exposed to high salinity. **A.** Western blot analysis of the abundance of RubisCO large subunit, PRK, LHCB1, LHCB2, LHCA1, PsbO, PsbA, NDH-H, PGR-L1, ATP synthase γ subunit, PSBS and FBN/CDSP34 in plants grown in standard conditions or exposed to high salinity treatments, 0.125 M NaCl or 0.1 M NaHCO_3_. **B.** Histogram representation showing the abundance of PsbO, PsbA and ATP synthase γ subunit in plants grown in standard conditions or exposed to high salinity treatments. Data are means ± SD of three or four values originating from independent experiments. * and **, significantly different from the WT value with P < 0.05 and P < 0.01, respectively (t-test). WT, wild type; D4, line co-suppressed for CDSP32; D10, line overexpressing CDSP32; DM19, line overexpressing CDSP32 active site mutant.

We then investigated the level of several thylakoid proteins involved in light collection, electron transfer, or protective mechanisms. The abundance of major proteins of light-harvesting complexes, LHCB1 and LHCB2, was very similar in the four potato lines in control and salt conditions. In contrast, in comparison with control conditions, the LHCA1 amount was increased upon NaCl watering, and noticeably decreased in plants treated with NaHCO_3_, particularly those overexpressing mutated CDSP32 (Fig. 7A and S9). Regarding PSII components, we noticed in control conditions a strong decrease (more than 50%) in the abundance of PsbO, one the three subunits of the PSII oxygen-evolving complex located in lumen, in CDSP32 over-expressing plants compared to WT (Fig. 7A lanes 1, 3 and 7B). Further, the protein level was also found to be substantially lower in this line and the one expressing mutated CDSP32 upon NaHCO_3_ treatment. Most interestingly, similar changes were observed for the D1 protein that together with D2 forms the reaction centre of PSII. Indeed, the abundance of D1 was significantly lower in CDSP32 over-expressing plants in control conditions and in plants expressing the mutated TRX upon NaHCO_3_ exposure (Fig. 7A lanes 1, 3 and 7B).

Zhang et al. (2021) proposed that CDSP32 could regulate photosynthetic cyclic electron transport. Therefore, we investigated the abundance of NDH-H and PGRL1 proteins, which are components of the two pathways operating around PSI (Rumeau et al. 2007; Hertle et al. 2013). Concerning NDH-H, a subunit of NAD(P)H dehydrogenase complex, we observed a strongly reduced protein level in all lines watered with NaCl compared to control and alkaline-saline conditions and no substantial genotype-dependent variation. Conversely, the abundance of PGRL1, which is involved in the cyclic pathway named proton gradient regulation route, was much higher in plants watered with NaCl, the protein being barely detected in plants grown in control conditions or treated with NaHCO_3_.

We previously reported that the ATP-synthase γ subunit interacts *in planta* with CDSP32 in plants overexpressing the mutant TRX form (Rey et al. 2005). In control conditions, we observed a lower amount of this subunit in plants overexpressing the non-mutated TRX than in WT (Fig. 7A lanes 1, 3 and 7B). Salt treatments resulted in an increased abundance of the γ subunit, with a lower level in plants overexpressing CDSP32 compared to WT upon exposure to NaHCO_3_ (Fig. 7B).

We analysed the abundance of PsbS, a photoprotective protein involved in dissipation of excess energy in conditions of high illumination (Li et al. 2000). We did not observe any significant change in the protein level in relation with CDSP32 expression level, but noticed a slightly higher protein abundance in all lines exposed to NaCl (Fig. 7A). Finally, we examined the abundance of fibrillins (FBN), a family of proteins possessing a lipid-binding motif and presumed among other functions to reduce photooxidative damage to PSII (Yang et al. 2006; Kim and Kim, 2022). We used the serum that we generated against FBN clade1-type in potato, initially named CDSP34 for Chloroplastic-Drought induced Stress Protein of 34 kDa (Pruvot et al. 1996). In control conditions, two FBNs were revealed at around 34 and 37 kDa, the latter form being more abundant in plants lacking CDSP32 or over-expressing the mutated TRX. Upon NaCl treatment, the FBN1 isoform was much more abundant in the four lines, consistently with our previous results (Pruvot et al. 1996). In contrast, watering with NaHCO_3_ resulted in a strong decrease of the abundance of the two isoforms in all lines.

### PSII function and activity of the electron transport chain

Our biochemical analyses revealed decreased abundance of the PsbO and D1 subunits in lines overexpressing CDSP32 in the absence of stress (Fig. 7). We performed fluorescence measurements to assess the effects of altered CDSP32 expression on the maximal PSII quantum yield (Fv/Fm) in dark-adapted leaves and the yield during the transition from darkness to moderate-low light (80 µmol photons m^-2^ s^-1^, Fig. 8A). Fv/Fm values were similar in all genotypes (∼0.79, Fig. 8C and Table S2), indicating a normal PSII function. The transient fluorescence increase in the first seconds of moderate-low illumination was instead significantly higher in the CDSP32 co-suppressed line (D4, Fig. 8B), although it then decreased to similar stationary levels as in the other genotypes during the following 4 minutes of illumination. This resulted in a lower PSII yield (Φ_PSII_) measured in D4 plants 4 seconds after the light onset (Fig. 8D and Table S2), but a normal Φ_PSII_ *ca.* 200 seconds after the light onset (Fig. 8E and Table S2). We observed an increased transient fluorescence in D4 also during the transition from darkness to moderate-high light (170 µmol photons m^-2^ s^-1^), although the difference was less pronounced because the increase was generally higher in all genotypes (Fig. S10A, B). In high light (750 µmol photons m^-2^ s^-1^), the initial fluorescence increase reached values close to Fm in all plants (Fig. S10C, D). The Non Photochemical Quenching (NPQ) of Fm’ induced by the medium-high and high lights was similar in all lines (Fig. S10A, C). At all light intensities, the fluorescence rise kinetics in the first hundreds of milliseconds of illumination were similar in all genotypes, suggesting a similar PSII antenna size.

**Figure 8.**
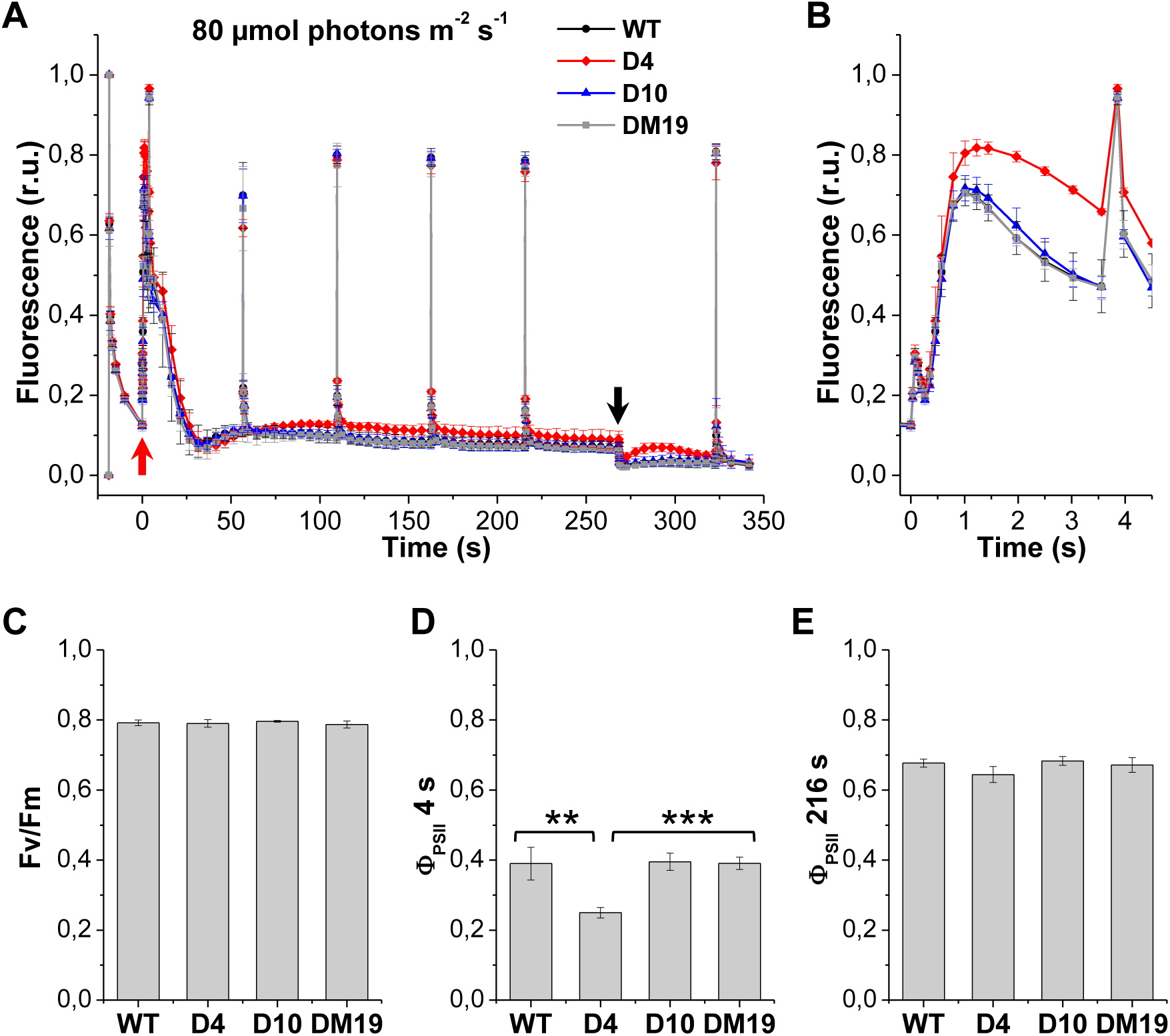
Chlorophyll fluorescence parameters in medium-low light. (A) Fluorescence induction curves recorded by applying to dark-adapted leaf fragments a multi-turnover saturating pulse followed by 4 minutes of actinic illumination at 80 µmol photons m^-2^ s^-1^ with superimposed saturating pulses. All curves are normalised on Fm-Fo. The red and black arrows indicate the onset (time 0) and offset of the actinic illumination. The averages ± S.D. of 3 biological replicates per genotype are shown. The first 4.5 seconds from actinic light onset of the curves are shown with an expanded time scale in panel (B), to better display the increased transient fluorescence in D4. (C), (D) and (E) Maximal PSII quantum yield (Fv/Fm), and PSII yield (Φ_PSII_) after 4 s and 216 s of actinic illumination, respectively, calculated from the curves in (A). All values represent the averages ± S.D. of 3 biological replicates and are reported in Table S2. The Φ_PSII_ 4 s in D4 is significantly lower than in the other genotypes (t-test, ** P ≤ 0.01, *** P ≤ 0.001).

In agreement with the fluorescence results, the PSII+PSI antenna size and total steady-state electron transport rates measured by ElectroChromic Shift (ECS) in function of light intensity were also similar in all genotypes (Fig. S11).

### Light-dependent activation of the ATP-synthase activity

Zimmer et al. (2021) have shown that, during the transition from darkness to low light, the reduction of CDSP32 follows kinetics similar to those of TRX f and of ATP-synthase γ subunit, and here we revealed a lower abundance of this subunit in lines overexpressing CDSP32 in control conditions (Fig. 7). Based on these results and on the *in planta* interaction between CDSP32 and the γ subunit (Rey et al. 2005), we investigated the redox-dependent activation of ATP-synthase in dark-adapted potato leaves.

The ElectroChromic Shift (ECS) measures the transthylakoid membrane potential (Δψ) generated by the photosynthetic electron and proton transport activity, whose dissipation mostly depends on the ATP-synthase activity. We thus measured the activity of the ATP-synthase based on the kinetics of ECS decay after a single-turnover flash, inducing charge separation, as previously done by Kramer and Crofts (1989). Since high Δψ levels can also induce an increase in ATP-synthase activity (Kramer and Crofts, 1989), we used sub-saturating flashes inducing charge separation in ∼55-59% of photosystems in all genotypes (Table S3).

We measured the flash-dependent ECS decay kinetics 2 minutes after different numbers of pre-flashes. At 2 minutes, the Δψ generated by the pre-flashes is dissipated, while the ATP-synthase reduced by the pre-flashes is not yet re-oxidised. In all genotypes, the ECS decay kinetics were slow in dark-adapted leaves, indicating an oxidised and thus inactive ATP-synthase, and fully accelerated after a train of 300 pre-flashes, that induced the reduction-dependent activation of the ATP-synthase. Of note, the acceleration for intermediate numbers of pre-flashes was delayed in the D4 line (Fig. 9A). Consequently, the number of charge separations required to achieve a 50% acceleration of the slow ECS decay phase was significantly higher in D4 than in the other genotypes (Fig. 9B, Fig. S12A, B and Table S4). In the DM19 line, expressing a mutated inactive form of CDSP32 in addition to the WT form, we observed a higher degree of variability in the acceleration of the ECS decay kinetics than in the other genotypes. For this reason, we present the results obtained in 5 DM19 biological replicates, instead of 3. Overall, the average number of charge separations required to achieve a 50% acceleration of the slow ECS decay phase in DM19 plants was still significantly lower than in D4 plants (Fig. 9B, Fig. S12B and Table S4).

**Figure 9.**
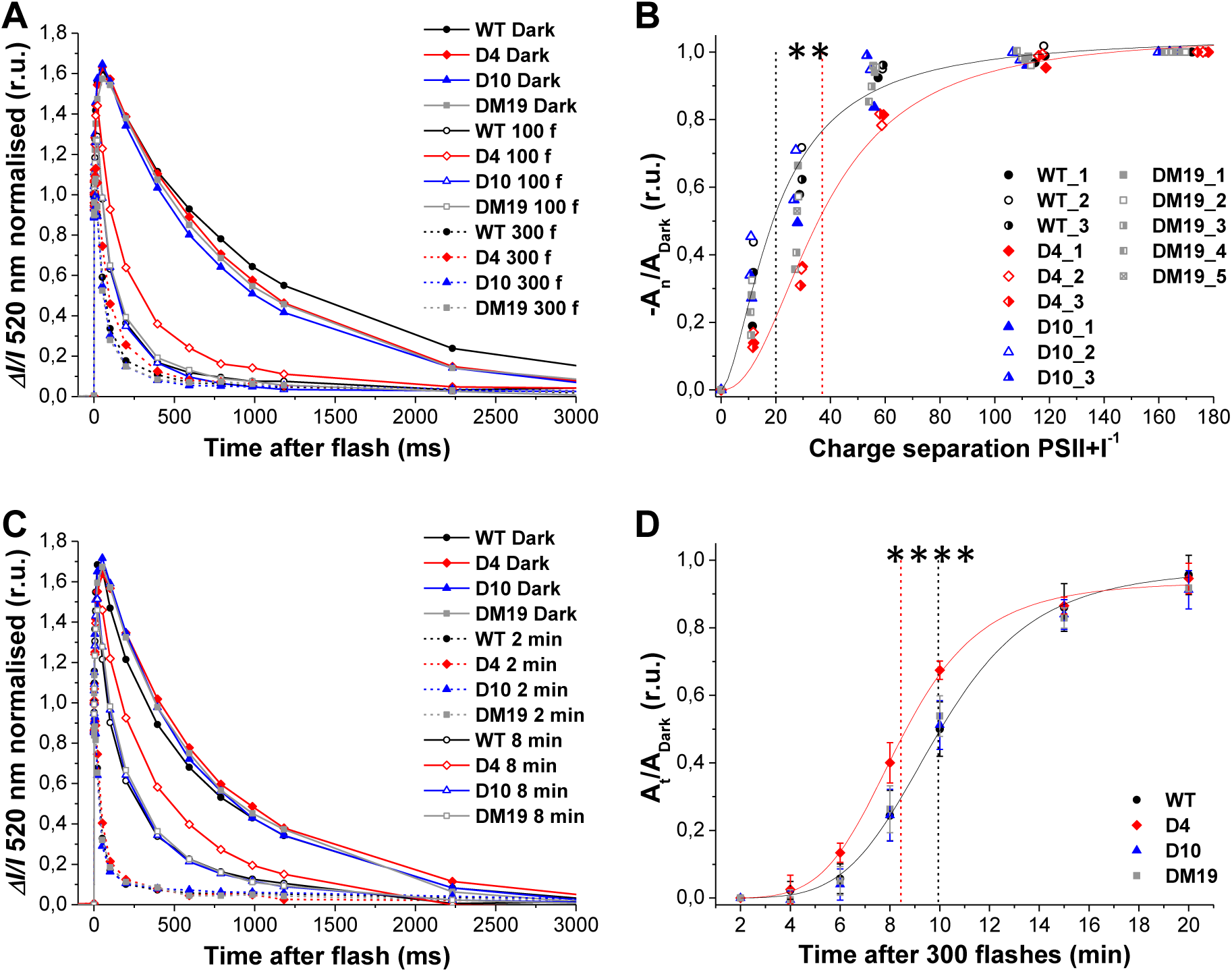
Redox-dependent regulation of the chloroplast ATP-synthase activity measured by ECS. (A) Representative flash-dependent ECS curves recorded in dark-adapted leaves (closed symbol, solid line), or 2 minutes after a train of 100 (open symbol, solid line) or 300 (closed symbol, dashed line) pre-flashes. (B) Loss in amplitude of the slow ECS decay phase induced by a variable number of pre-flashes applied to dark-adapted leaves (–A_n_/A_Dark_), plotted in function of the number of charge separations PSII+PSI^-1^ generated by the pre-flashes (averages ± S.D. of 3 biological replicates for WT, D4 and D10, 5 replicates for DM19). The black and red solid lines represent the Hill fit of the WT and D4 datasets, respectively, and the vertical dashed lines indicate the number of charge separations required to achieve a 50% acceleration of the slow ECS decay phase in the two samples. This number is significantly higher in D4 (t-test, ** P ≤ 0.01). The fits the D10 and DM19 datasets are shown in Fig. S12A and B, all fit results are reported in Table S4. (C) Representative flash-dependent ECS curves recorded in dark-adapted leaves (closed symbol, solid line), or 2 minutes (closed symbol, dashed line) or 8 minutes (open symbol, solid line) after a train of 300 pre-flashes. (D) Recovery in amplitude of the slow ECS decay phase during a variable dark incubation time after a train of 300 pre-flashes applied to dark-adapted leaves (A_t_/A_Dark_), plotted in function of the dark incubation time (averages ± S.D. of 4 biological replicates per genotype). The black and red solid lines represent the Hill fit of the WT and D4 datasets, respectively, and the vertical dashed lines indicate the time of dark incubation required to achieve a 50% recovery of the slow ECS decay phase in the two samples. This time is significantly shorter in D4 (t-test, **** P ≤ 0.0001). The fits the D10 and DM19 datasets are shown in Fig. S12C and D, all fit results are reported in Table S5.

We then investigated the ATP-synthase de-activation in darkness by measuring the flash-dependent ECS decay kinetics at different times after a train of 300 pre-flashes. While the ECS decay kinetics were equally fast in all genotypes 2 minutes after the pre-flashes, their subsequent deceleration was faster in the D4 line (Fig. 9C). Consequently, the dark time required to achieve 50% recovery of the slow ECS decay phase was significantly shorter in the D4 line (Fig. 9D, Fig. S12C, D and Table S5).

## Discussion

### Presence and specificities of CDSP32 in the green lineage

The atypical CDSP32 TRX exhibits very unique features with the presence of two TRX-fold domains in tandem and an atypical HCGPC redox active site motif in only one of them. The search for homologues among living organisms revealed the presence of this TRX type in many classes of green plants, with a high degree of conservation particularly in Angiosperms, and to a lesser degree in Pteridophytes. Related proteins showing divergence in the active site motif are present in Bryophytes, Charophytes and Chlorophytes, with some Algae sequences carrying only one Cys in the active site (Fig. 1), implying distinct biochemical functions in these organisms. Most importantly, no related protein was identified in Cyanobacteria, indicating a non-prokaryotic origin of the gene. Very surprisingly, a few bacteria genomes harbour sequences highly homologous to Angiosperm CDSP32 (Fig. S4). This unexpected ranking of sequence similarity is a strong argument for horizontal gene transfer from the plant kingdom in the commensal microbiota (Koonin et al. 2001). The remarkable conservation of CDSP32 in the main classes of vascular plants suggests that this atypical TRX fulfils specific physiological functions that cannot be performed as efficiently by other plastidial TRXs, despite their remarkable diversity in higher plants (Meyer et al. 2012).

### Participation of the CDSP32 TRX in redox homeostasis maintenance upon salt stress

The first data gained on transgenic potato plants highlighted the participation of CDSP32 in responses to oxidative stress (Broin et al. 2002). We proposed that the TRX fulfils an antioxidant role allowing the maintenance of plastidial redox homeostasis upon environmental constraints. TRXs are now recognized as essential components in redox and ROS-related transduction pathways allowing integration of multiple signals generated by climatic constraints (Vieira dos Santos and Rey, 2006; Mittler et al. 2022). This is illustrated by the participation of apoplastic TRX h, mitochondrial TRX o, and plastidial TRXs m in responses to salt or light constraints in rice and Arabidopsis (Zhang et al. 2011; Calderon et al. 2018; Serrato et al. 2021). However, while we observed an increased CDSP32 abundance in WT potato leaves upon various NaCl treatments (Fig. 5A; Pruvot et al. 1996), no phenotype was noticed in plants modified in *CDSP32* expression under such a salt exposure (Fig. S7). This might be due to insufficient severity of the treatment applied since WT plants did not exhibit any damage. Indeed, potato is classified as a moderately salt-sensitive crop (Hütsch et al. 2018; Chourasia et al. 2021). We cannot exclude that CDSP32 participates in responses to higher harmful NaCl concentrations that affect redox homeostasis (Munns and Tester, 2008). Consistently, we observed upon high salinity a decreased CDSP32 level in *Arabidopsis* while its abundance was unaffected in a halophyte related-species (Mrah et al. 2007).

Here we unveil the participation of the TRX in responses to NaHCO_3_ treatment, which generates alkaline stress in addition to ionic and osmotic stresses compared to exposure to neutral salts. Potato plants developed much more severe damage when watered for the same period with 0.1 M NaHCO_3_ than with 0.125 M NaCl, showing that the phenotype does not originate from sodium toxicity. Furthermore, plants overexpressing a mutated form were more susceptible to this treatment as evidenced by growth and chlorophyll content measurements (Fig. 4). Plant adaptation to saline-alkaline stress has been only recently investigated (Cao et al. 2022; Rao et al. 2023), and involves common responses with neutral salt stress like hormone signalling, accumulation of osmolytes and regulation of ion homeostasis, but also more specific responses to limit pH modification within plant cells (Cao et al. 2022). Saline-alkaline conditions strongly impair ROS production levels and cell redox homeostasis (Rao et al. 2023). Consistently, we observed that exposure to NaHCO_3_, unlike to NaCl, results in strongly reduced abundance of cytosolic APX in potato plants (Fig. 6A). Elevated ROS-scavenging capacity and increased glutathione content have been shown to be associated to enhanced resistance to saline-alkaline stress in poplar (Wang et al. 2020) and alfalfa (Sun et al. 2020), respectively. Our data on CDSP32-modified lines reveal the participation of a TRX-based mechanism in the responses to the redox changes induced by alkaline stress. CDSP32 could be involved in a pathway regulating the ROS-scavenging capacity since the three lines modified for its expression displayed a reduced CAT abundance. In addition, we noticed a significantly higher abundance of Cu/Zn SOD in the line overexpressing the TRX. In other respects, we observed substantial changes in the abundance and/or redox status of CDSP32 partners in saline-alkaline stress conditions. For instance, compared to WT, potato plants lacking the TRX or overexpressing its non-functional form displayed a much higher level of over-oxidized 2-Cys PRX, and the three modified lines exhibited higher abundances of plastidial MSRBs upon NaHCO_3_ exposure (Fig. 5). PRXs and MSRs participate in ROS scavenging pathways or in signaling mechanisms (Liebthal et al. 2018; Rey and Tarrago, 2018). Interestingly, overexpression of 2-Cys PRX was recently reported to confer tolerance to NaHCO_3_, that was associated to maintained photosynthetic activity and increased ROS scavenging capacity (Wang et al. 2023). In other respects, Sun et al. (2016) revealed interaction of one cytosolic methionine sulfoxide reductase B with a Ca^2+^-kinase in soybean, leading to activation of ROS-signalling under alkaline stress. Altogether, these findings prompt us to propose that CDSP32 and its partners participate in plant responses to saline-alkaline stress, and could prevent damage in components of photosynthetic electron transfer and chlorophyll biosynthesis pathways.

### Regulation by CDSP32 of photosynthesis and ATP-synthase activation

#### Abundance of components of the photosynthetic apparatus

Up to now and in comparison with typical TRXs f or m, the CDSP32 TRX has been considered to be mainly involved in the maintenance of plastidial redox homeostasis upon stress conditions through interaction with other thiol-reductases such as PRXs and MSRs (Rey et al. 2005). In this work, we provide several lines of evidence showing that the TRX is involved in the regulation of photosynthetic metabolism at several levels in the absence of stress. Indeed, the DM19 line over-expressing mutated CDSP32 exhibits a reduced chlorophyll content (Fig. 3D), revealing that the presence of a high proportion of inactive CDSP32 affects chlorophyll metabolism and/or biosynthesis. The activity of several enzymes involved in chlorophyll biosynthesis is redox-regulated by various plastidial TRX types including NTRC in relation with 2-Cys PRX (Stenbaek et al. 2008; Da et al. 2017; Richter et al. 2018). The phenotype trait of DM19 could result from irreversible trapping by the mutated TRX of redox-sensitive enzymes involved in chlorophyll metabolism or from the modified redox status of 2-Cys PRX observed in this line (Fig. 5A). In control conditions, the PSII D1 and PsbO subunits accumulated to significantly lower levels in plants overexpressing CDSP32 than in WT (Fig. 7). Of note, Wang et al. (2013) reported substantially reduced levels of these two PSII subunits in Arabidopsis plants lacking three TRX m isoforms, and also showed redox-based interaction between the D1 subunit and TRXs m. PsbO also interacts with TRX (Montrichard et al. 2009) and thus could be possibly redox-regulated through sensitive cysteines, which would be reduced via a NTRC-dependent pathway providing electrons to this PSII subunit located in the lumen (Ameztoy et al. 2019). A lower abundance of D1 and PsbO subunits could result from impaired expression or increased degradation due to redox modifications, and might be associated with subsequent impairment of PSII function. Nonetheless, the PSII maximum quantum yield (Fv/Fm, Fig. 8C) is normal in the overexpression line, indicating that assembled PSII complexes properly function. From the protein analyses in Fig. 7, we did not observe any difference regarding PSI and PSII light harvesting antennas (LHCA1 and LHCB1 and 2, respectively) in any of the genotypes grown upon standard conditions. This agrees with the fact that the functional antenna size of PSII measured by fluorescence (Fig. 8B and S10B, D) and of PSII+PSI measured by ECS (Fig. S11B) are similar in all lines. Taken together, these data indicate that PSII function and antenna size are independent on CDSP32 abundance.

On the other hand, the fluorescence induction curves show that electron transport downstream of PSII is transiently impaired in the line lacking CDSP32 during dark/light transitions (Fig. 8B, D). This suggests a delay in the early stages of photosynthesis activation. Once the activation is completed, the activity of the electron transport chain is independent of *CDSP32* expression, as indicated by the similar levels of stationary Φ_PSII_ and NPQ (which depends on the lumen acidification induced by electron transport) reached in all genotypes (Fig. 8 and S10). This is further confirmed by the similar steady-state electron transport rates measured by ECS in function of light intensity (Fig. S11).

#### Redox-dependent regulation of ATP-synthase activity during dark/light and light/dark transitions

Zimmer et al. (2021) have shown that, during the transition from darkness to low light (40 µmol photons m^-2^ s^-1^), the activation of chloroplast ATP-synthase via the reduction of its γ subunit is completed within the first 30 seconds of illumination, while the redox-dependent activation of the Calvin-Benson cycle enzymes is slower (a couple of minutes). The reduction of CDSP32 and TRX f also peaks at 30 seconds of illumination. This agrees with the proposal made by Kramer and Crofts (1989) that the ATP-synthase activation, which only requires very low light intensities, depends on the redox equilibrium between the γ subunit and TRXs, and should not be limiting for photosynthesis in normal conditions. In our induction curves in medium-low light (Fig. 8), we can attribute the initial transient increase of fluorescence to an over-reduction of the electron transport chain caused by an inactive CBC. This fluorescence transient is higher in the absence of CDSP32 at few seconds after the light onset, but then decreased with similar kinetics in all genotypes during the first minute of illumination. This suggests a similar activation of Calvin-Benson cycle enzymes in all genotypes, with the fluorescence phenotype in the D4 line being due to an impairment in a process that occurs over a shorter time-scale. Based on the previously detected *in planta* interaction between CDSP32 and the γ subunit (Rey et al. 2005), we hypothesize this process to be the activation of the ATP-synthase. Indeed, our ECS measurements show that the light-dependent activation of the ATP-synthase still occurs in the absence of CDSP32 (D4 plants), but requires more charge separations in PSII and PSI than in the WT, and therefore a higher reduction level of the stroma (Fig. 9B). Concomitantly, the dark reoxidation of the ATP-synthase after illumination is faster in the absence of CDSP32 (Fig. 9D). When the mutated CDSP32 is expressed in addition to the active form (DM19), the kinetics of ATP-synthase activation remain on average faster than in absence of the TRX, but present a higher degree of variability between biological replicates (Fig. 9B and Fig. S12B), possibly reflecting a variability in the relative accumulation levels of the two CDSP32 forms. Overexpression of the WT form of CDSP32, instead, does not cause any significant difference in the kinetics of ATP-synthase activation and de-activation. Based on these results, we propose that CDSP32 is responsible for the early stages of redox activation of the ATP-synthase (via the γ subunit) during dark/light transition. Additionally, CDSP32 seems to mitigate ATP-synthase reoxidation during light/dark transition, possibly acting as a storage of reducing power due to its high abundance in the chloroplast. In D4 plants lacking CDSP32, this redox regulation could be performed by another thioredoxin, but with delayed kinetics. Zimmer et al. (2021) have shown that TRX f is reduced with very similar kinetics to both CDSP32 and ATP-synthase γ subunit during the initial phases of illumination, so this canonical TRX could fulfil the CDSP32 function in co-suppressed plants. In DM19 plants, the mutated CDSP32 could prevent a fraction of the ATP-synthase to be activated during the dark/light transition, in competition with the WT form or TRX f, by irreversible trapping of the γ subunit.

Assuming that CDSP32 and TRX f are in thermodynamic equilibrium (as suggested by Zimmer et al. 2021), we can use the data in Fig. 9B and D to calculate the equilibrium constant (*K*) between the two TRXs, as detailed in the Supplementary Information. The calculated equilibrium constant is of *ca.* 0.3 for the flash-dependent activation data and of *ca.* 0.5 for the dark-dependent de-activation data (Fig. 10A and B, respectively). Based on these values, the Em of CDSP32 should be between 16 and 9 mV higher that the Em of the putative substitute TRX, TRX f. This is very much in line with the ΔEm that would be expected based on the published Em values of the two proteins (*ca.* −290 mV for TRX f and −280 mV for CDSP32, as reported in Zimmer et al. 2021). Our results suggest that CDSP32, thanks to its slightly higher Em, could ensure a fast redox-dependent activation of the ATP-synthase when plants are exposed to low light, and regulate the kinetics of its de-activation in darkness.

**Figure 10.**
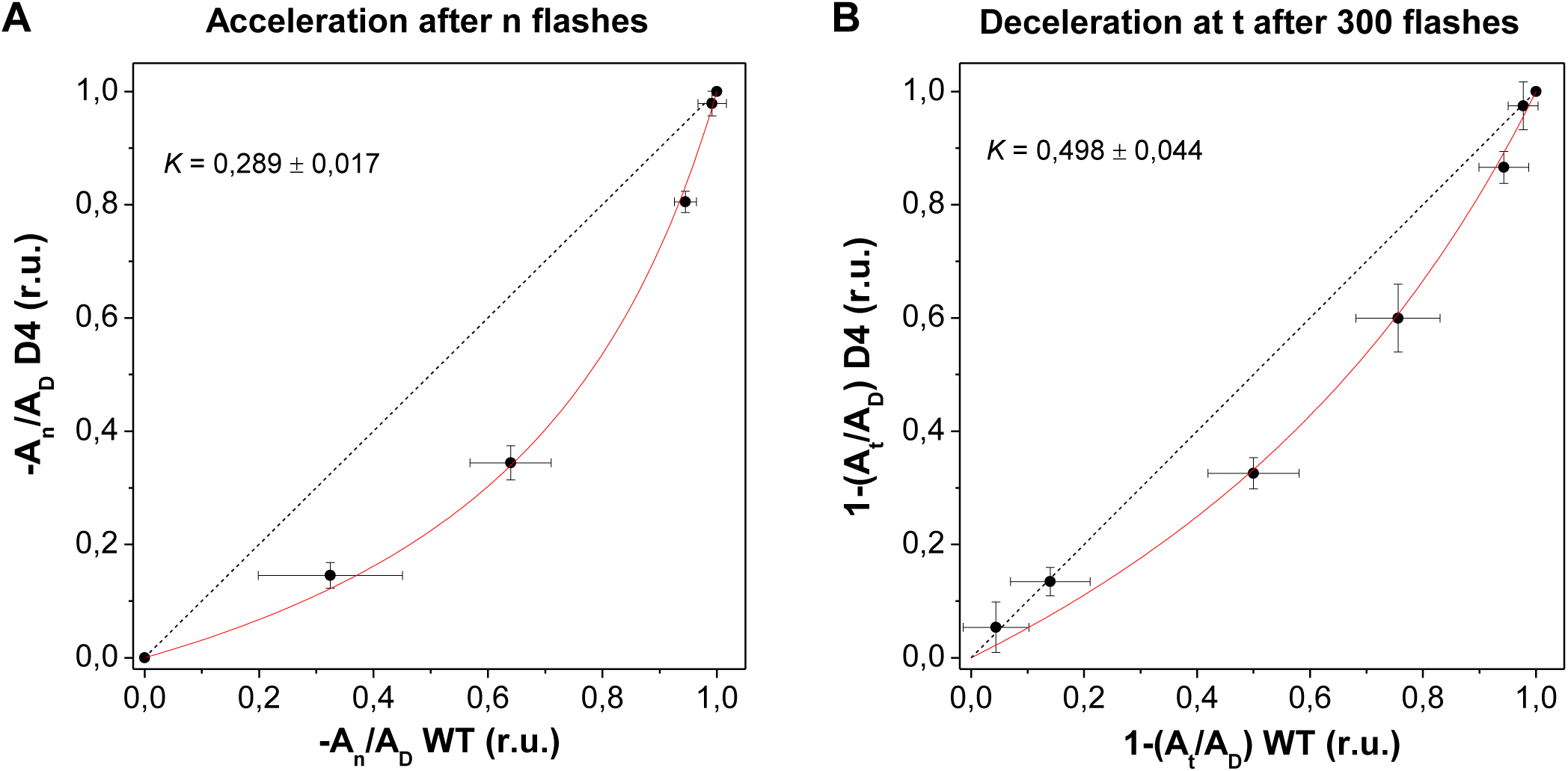
Comparison of the redox-dependent regulation of the chloroplast ATP-synthase activity in WT and D4 plants. (A) Loss in amplitude of the slow ECS decay phase induced by a variable number of pre-flashes applied to dark-adapted leaves (–A_n_/A_Dark_) in D4 versus WT plants (averages ± S.D. of 3 biological replicates per genotype, using the data in Fig. 9B). (B) Inverse of the recovery in amplitude of the slow ECS decay phase during a variable dark incubation time after a train of 300 pre-flashes applied to dark-adapted leaves (A_t_/A_Dark_) in D4 versus WT plants (averages ± S.D. of 4 biological replicates per genotype, using the data in Fig. 9D). In both panels, data were fitted with the equation 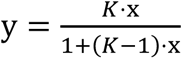 (see Supplementary Information for details). The *K* ± S.E.M. values obtained by the fit are reported on each graph.

The ATP-synthase has been identified as one of the first photosynthetic components regulated by the TRX system (Mc Kinney et al. 1978) via the control of the redox status of two Cys residues situated in a 9-residue loop domain of the γ subunit (called the redox loop) (Akiyama et al. 2023). Various plastidial TRX systems depending on NTRC or FTR (via TRXs f and m) have been reported to modulate the activity of the ATP-synthase complex (Yoshida et al. 2022; Diaz et al. 2020; Sekiguchi et al. 2020). Rapid reduction of ATP-synthase γ subunit has been noticed under low-light conditions that do not lead to reduction of CBC enzymes, revealing differential dependency on linear electron transfer between this subunit and enzymes like SBPase or FBPase (Hisabori et al. 2013; Yoshida et al. 2014). These authors hypothesized that a closer proximity of the ATP-synthase complex with the PSI/Fd/FTR/TRX system could explain its more efficient reduction compared to stromal enzymes. Of note, the NTRC system has been proposed to be a key activator of the ATP-synthase under low irradiance conditions (Nikkanen et al. 2016). Our data reveal that plants lacking CDSP32 display reduced activation of ATP-synthase during transition from dark to low light, indicating that other plastidial TRXs cannot compensate as efficiently the absence of the TRX. This likely originates from specific intrinsic properties of this atypical TRX such as a slightly higher E_m_, or the more negative electrostatic surface charge compared to TRX f (Fig. 2). Based on the present results and the previously detected *in planta* interaction between *S. tuberosum* CDSP32 and the ATP-synthase γ subunit (Rey et al. 2005), the 3D structure of the complex formed between CDSP32 and the γ subunit of *S. tuberosum* ATP-synthase was modelled using AlphaFold2 to see how these two proteins could interact. Chloroplast ATP-synthase is a macromolecular machine made of 26 protein subunits, 17 of them being embedded in the membrane and the regulatory γ subunit being part of the central stalk as shown in the cryo-EM structure of *Spinacia oleracea* ATP-synthase (Hahn et al. 2018). The AlphaFold2 3D structure of *S. tuberosum* ATP-synthase γ subunit alone shows the same folding as the spinach γ subunit (PDB 6FKF) (83.4 % identity between these two γ-subunits) with two long α-helices in a tight coiled coil that forms the rotor shaft and a L-shaped structure containing the redox loop and a β-hairpin (Fig. 11A, Fig. S13A-B). In the ATP-synthase γ subunit-CDSP32 complex, the γ subunit redox loop and the redox active motif of CDSP32 TRX_D2 are in close proximity with a distance of 4Å between the S of the catalytic Cys of each partner, showing that CDSP32 can reduce the disulphide bridge of the γ subunit redox loop (Fig. 11B, Fig. S14). The superposition of the *S. tuberosum* γ subunit-CDSP32 complex on the 3D structure of *S. oleracea* ATP-synthase shows that CDSP32 TRX has sufficient space to interact with the γ subunit without steric hindrance (Fig 11B). The spinach ATP-synthase γ subunit-CDSP32 complex was also modelled using Afphafold2 and shows the same result (data not shown). It has to be noticed that the L-shaped structure present in the γ subunit alone is no longer structured in the ATP-synthase γ subunit-CDSP32 complex (Fig. 11). These changes are in agreement with the recent proposed model of redox regulation of the ATP synthesis by cooperative function of the redox loop and the β-hairpin (Akiyama et al. 2023).

**Figure 11.**
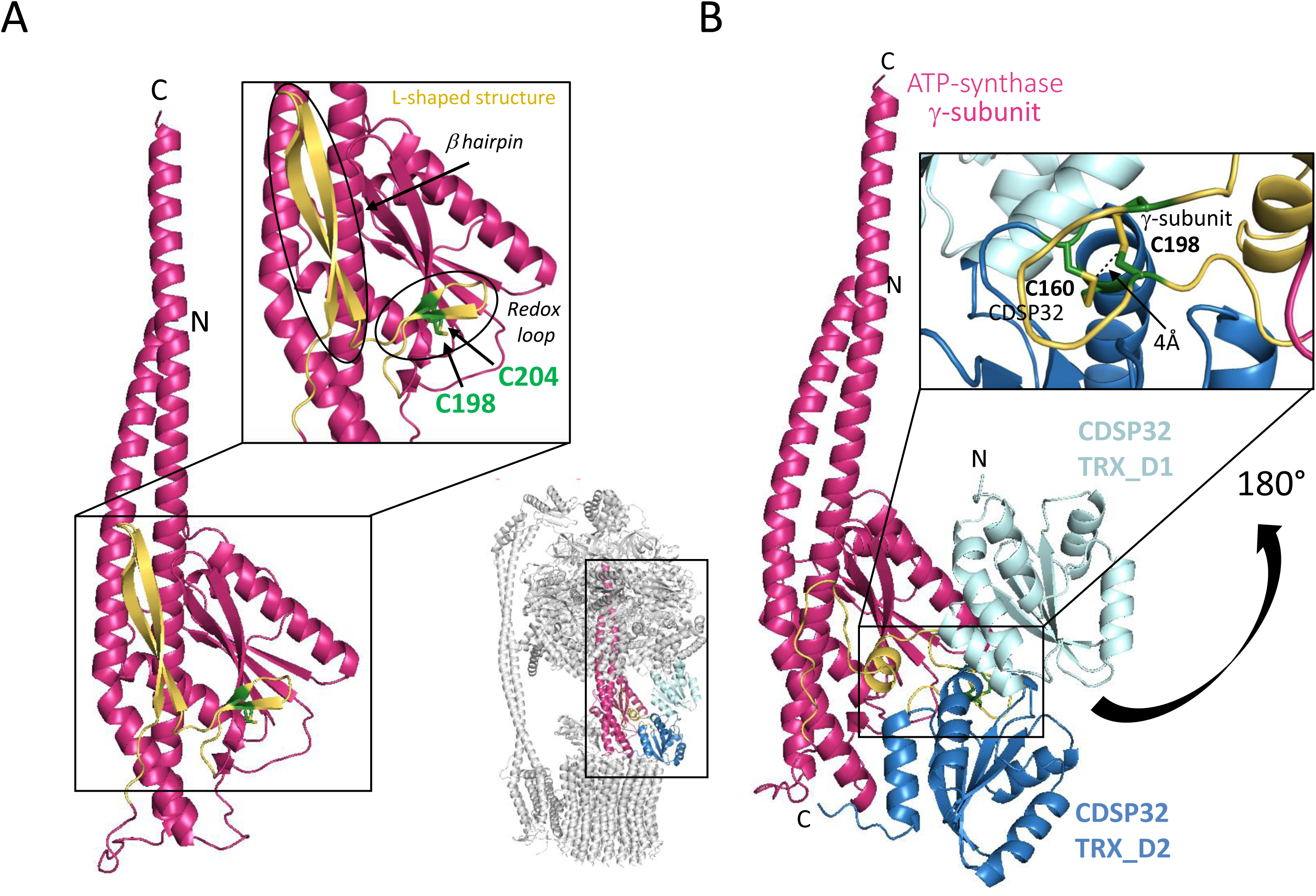
*S. tuberosum* CDSP32 in complex with ATP synthase γ subunit. (A) AlphaFold2 3D structure of *S. tuberosum* ATP synthase γ subunit alone showing the L-shaped structure (yellow) containing the redox loop and the β-hairpin, (B) Complex between CDSP32 and ATP synthase γ subunit generated by AlphaFold2 showing a close proximity between the γ subunit redox loop and the CDSP32 redox active motif, with a close proximity (4Å) of the two catalytic Cys residues of each partner. Superposition of the *S. tuberosum* CDSP32-ATP synthase γ subunit complex onto the spinach ATP-synthase (PDB 6FKF) is shown aside.

Modelisation of *S. tuberosum* TRX f in complex with the ATP-synthase γ subunit was also generated using AlphaFold2. In this complex, the redox loop of the γ subunit and the redox active motif of TRX f are not close, with a distance of 30Å between the S of the catalytic Cys of each partner (Fig. S15). This indicates that it is probably more difficult for TRX f to interact with the γ subunit. Indeed, in the γ subunit-CDSP32 complex, the interaction zone of CDSP32 is located almost exclusively in TRX_D2 (Fig. S14D) in a region showing different electrostatic properties with those of TRX f (Fig. 2B). Of note, the complex generated by AlphaFold2 between the γ subunit and CDSP32 TRX-D2 only shows the same position for TRX-D2 as in the complex involving the whole CDSP32 indicating that CDSP32 TRX_D1 might not be crucial for the interaction with the ATP-synthase γ subunit (data not shown). These data raise the question of the role of this N-terminal domain, which is unique to CDSP32. Other large TRXs carry domains with known functions such as protein disulphide isomerase activity or interaction with heat shock proteins (Meyer et al. 2012). Taking into account the beneficial stabilizing properties of the TRX-fold, which are widely used for producing recombinant proteins, TRX-D1 might help stabilizing and/or positioning the CDSP32 TRX within the ATP-synthase complex.

It is interesting to note that both CDSP32 and the region of the ATP-synthase γ subunit containing the two cysteines that are the targets of the redox regulation are present in plants and green algae, but absent in Cyanobacteria (Figs. 1, S16; Akiyama et al. 2023). It is tempting to speculate that the two partners could have evolved together after the endosymbiotic event, from which the chloroplast originated, and that CDSP32 could have then been lost later in some organisms (Fig. 1). Indeed, a redox fine-tuning of the ATP-synthase activity during light transitions makes sense in the chloroplast, where the stroma is relatively isolated from the surrounding cellular environment, and its redox poise closely reflects the photosynthetic electron transport activity. In Cyanobacteria, instead, the thylakoid membranes are embedded in the cytoplasm, whose redox poise does not change rapidly in function of the sole photosynthetic electron transport activity. Sugar catabolism reduces the cytosolic NADPH pool also in darkness to sustain the activity of the respiratory electron transport chain (Mi et al. 1994) that is also located in the thylakoids (Mullineaux, 2013). The reducing conditions maintain the ATP-synthase active in darkness (Viola et al. 2019), to produce ATP using the pmf generated by respiration. When extending the search for CDSP32 homologues in other eukaryotic lineages, we could not find any in red Algae and in organisms whose chloroplast derives from them, such as diatoms and coccolithophores. Interestingly, the ATP-synthase γ subunit of these organisms also lacks the two cysteines that are the targets of the redox regulation in plants and green algae (Fig. S16). This supports the idea of a possible evolutionary correlation between the presence of CDSP32 and the redox fine-tuning of the ATP-synthase activity, that seems to be specific to the green eukaryotic lineage. The absence of this fine-tuning in red Algae and derivatives might reflect a closer metabolic interaction between chloroplasts and mitochondria, as characterised in diatoms (Bailleul et al. 2015), that could partially uncouple the redox poise of the stroma from photosynthetic electron transport.

## Conclusions

Altogether, our findings reveal that the CDSP32 TRX fulfils crucial functions both in the plastidial antioxidant network by providing electrons to various thiol reductases upon environmental constraints and in the direct regulation of photosynthetic metabolism in the absence of stress via the control of ATP-synthase activity. Of note, this TRX is one of the most highly expressed in photosynthetic cells, and its synthesis is strongly triggered upon stress conditions (Rey et al. 1998; Broin et al. 2000; Belin et al. 2015). This further argues for an essential function of CDSP32 in the maintenance of chloroplast redox homeostasis and direct regulation of photosynthetic activity in relation to environmental conditions. CDSP32 interacts with the 2-Cys PRX, a major plastidial thiol peroxidase acting as a redox sensor of H_2_O_2_, the concentration of which is increased upon stress conditions (Vogelsang and Dietz, 2020). H_2_O_2_ reacts with 2-Cys PRXs (Dietz, 2008), which in turn can oxidize TRXs and deactivate photosynthetic enzymes (Yoshida et al. 2018). H_2_O_2_ would thus fulfil, in addition to an electron sink role, a regulatory function via thiol reductases under stress conditions (Driever and Baker, 2011). In addition to its role in the regulation of ATP-synthase activity during light/dark and dark/light transitions in the absence of stress, CDSP32 could thus regulate the activity of the photosynthetic machinery under stress conditions leading to ROS production.

Salt-induced osmotic stress conditions results in increased ATP-synthase abundance particularly upon NaCl treatment (Fig. 7A, B) Interestingly, Kohzuma et al. (2009) reported a substantially reduced abundance of this subunit in watermelon plants subjected to drought. Thus, such stress conditions are likely to result in impaired activity of this complex. Kohzuma et al. (2009) proposed that ATP-synthase represents a key point allowing proper photosynthetic proton circuit during acclimation of plant to long-term environmental constraints. It is tempting to speculate that the higher CDSP32 abundance observed in drought-stressed potato plants (Rey et al. 1998) could be a mechanism to maintain ATP-synthase activation despite a reduced amount of this complex. In other respects, all potato lines exhibited strongly reduced growth and decreased chlorophyll content when exposed to NaHCO_3_ (Fig. 4). This could originate from pH alteration within plastidial sub-compartments due to saline-alkaline stress, and subsequent impaired functioning of ATP-synthase. Of note, the line expressing the CDSP32 mutant form was found to be more sensitive than WT to NaHCO_3_, possibly due to more severe disturbance of the ATP-synthase machinery resulting from irreversible trapping of the γ subunit (Rey et al. 2005).

## Funding

This work was funded by a transverse project of the BIAM Institute and by Agence Nationale de la Recherche, Grant Award Number: ANR-23-CE20-0009.

## Acknowledgements

We are very grateful to Séverine Boiry and the “Phytotec” platform (CEA, DRF, BIAM) for technical assistance with growth chambers. We thank Dr Eevi Rintamäki (University of Turku, Finland) for providing the serum raised against Arabidopsis NTRC.

